# Ecological diversification without genomic reorganization: repeat dynamics and regulatory evolution in *Metarhizium robertsii*

**DOI:** 10.64898/2026.04.30.721964

**Authors:** Huiyu Sheng, Raymond J St. Leger

**Affiliations:** Department of Entomology, University of Maryland, College Park, Maryland, United States of America

**Keywords:** *Metarhizium robertsii* phylogenomics, evolutionary trajectory, metabolic flexibility, multifunctional ecological roles, repeat-induced point mutation, transposable elements

## Abstract

How can closely related organisms occupy distinct ecological niches without extensive genomic reorganization? We addressed this question by comparing eight strains of the entomopathogenic fungus *Metarhizium robertsii* that span a shallow phylogenetic gradient yet exhibit striking differences in virulence, plant associations, and metabolic capabilities. Despite nucleotide identity exceeding 98.5% and extensive macrosynteny, these strains showed pronounced variation in repeat-rich genomic regions: synteny gaps were twofold enriched near chromosome ends, transposable element loads varied from less than 3% to over 13% across strains diverging within ∼0.3 million years, and repeat-induced point mutation signatures tracked TE activity rather than phylogenetic distance. Notably, repeat expansion occurs through diverse spatial mechanisms rather than being uniformly concentrated in chromosome-terminal regions. In contrast, core functional repertoires, proteases, carbohydrate-active enzymes, developmental regulators, and most secondary metabolite biosynthetic genes, were highly conserved, with phenotypic differences in virulence and secondary metabolism arising primarily from regulatory divergence and local structural variation rather than gene presence or absence. Metabolic specialization similarly reflected functional repurposing within a conserved enzymatic framework: loss of processive cellulases alongside enrichment of oxidative auxiliary activity enzymes produced a profile convergent with brown-rot fungi but adapted for insect-associated and rhizosphere niches. These results support a hierarchical model of ecological diversification in which diverse repeat expansion mechanisms, TE dynamics, and regulatory innovation operate as coupled axes of localized genomic change within an otherwise constrained genomic framework, reconciling rapid niche differentiation with strong structural conservation and suggesting a general mechanism for intraspecific adaptation in complex eukaryotic microbes.

## Introduction

Understanding how ecological divergence arises over short evolutionary timescales remains a central challenge in evolutionary biology. Closely related organisms frequently exhibit substantial differences in host association, metabolism, and life-history strategies despite sharing highly conserved genomes. This apparent paradox raises a fundamental question: does ecological diversification require genome-wide innovation, or can it proceed through targeted modification of genomic and regulatory frameworks? Recent work increasingly suggests that phenotypic divergence may arise not only from changes in gene content, but also through evolution of regulatory networks, genome defense systems, and post-transcriptional control mechanisms, including small RNA–mediated processes(Verta and Jacobs 2024). Yet distinguishing between these alternative models requires high-resolution comparative analyses across closely related lineages, something that has proven challenging to achieve in most biological systems.

Fungal entomopathogens provide an exceptional system for addressing this question. These organisms navigate complex ecological challenges, requiring specialized mechanisms to breach insect cuticular defenses, survive host immune responses, and often establish beneficial relationships with plants as endophytes or rhizosphere associates. Comparative genomic analyses across distantly related taxa, including *Beauveria bassiana*, *Ophiocordyceps*, and members of the Hypocreales, have revealed repeated expansions of gene families associated with cuticle degradation, toxin production, and host interactions(Gao et al. 2011; Zheng et al. 2011; Xiao et al. 2012; Hu et al. 2013; Agrawal et al. 2015; Wichadakul et al. 2015; Agrawal et al. 2016). These findings suggest evolutionary convergence on a shared “entomopathogenic toolkit”(St. Leger and Wang 2020). However, comparisons across deeply divergent taxa are inherently limited in their ability to resolve evolutionary processes operating over short timescales, such as recent niche shifts, changes in virulence profiles, or transitions in host association patterns. To understand the genomic basis of rapid ecological differentiation, we need high-resolution analyses of closely related strains that exhibit clear phenotypic differences.

*Metarhizium robertsii* serves as an experimentally tractable model for investigating genome evolution across shallow phylogenetic scales. Prior phylogenomic and phenotypic analyses of eight strains revealed that variation in insect pathogenicity, plant endophytism, and metabolic profiles closely mirrors phylogenetic relationships (Sheng and St. Leger 2026). Early diverged strains (Mr1046, Mr727, and Mr1120) display slower germination, reduced virulence, and prolific sporulation—characteristics resembling those of early-evolved, narrow host range specialist *Metarhizium* species such as *M. acridum* (St. Leger and Wang 2020). In contrast, more recently diverged strains (Mr2547, Mr1878, Mr23, Mr1-16, and Mr2575) tend to be toxigenic, exhibit rapid germination, enhanced virulence, and stronger plant root associations (Sheng and St. Leger 2026). These findings suggest that ecological diversification within *M. robertsii* has occurred without extensive phylogenetic separation, providing an ideal framework to dissect the genomic mechanisms driving niche expansion.

Here, we test two competing models of ecological divergence within a species complex. Under a genome-wide innovation model, ecological differentiation should be accompanied by substantial changes in gene content, genome structure, and functional repertoires. Alternatively, a localized innovation model predicts divergence involving repeat-rich genomic regions and regulatory layers, including transposable element dynamics, genome defense processes such as RIP, transcriptional networks, and small RNA–mediated regulation. Critically, the localized innovation model predicts that divergence will occur within a shared set of repeat-rich genomic loci across strains, rather than through acquisition of lineage-specific genomic compartments. This distinction allows us to test whether genomic diversification in *M. robertsii* reflects evolutionary reuse of common dynamic regions or independent acquisition of novel genomic domains.

To distinguish between these alternative models, we performed a comprehensive comparative genomic analysis of eight *M. robertsii* strains representing early- and recently diverged lineages with distinct ecological traits. We integrated phylogenomics, genome architecture analysis, functional annotation, and experimental validation of metabolic activity to assess whether ecological diversification is driven by large-scale genomic restructuring or by localized evolutionary processes. Our results demonstrate that despite extensive conservation of genome structure and gene content, substantial ecological divergence has emerged through modifications involving dynamic genomic regions, though the spatial distribution of these modifications varies among strains. These findings provide a mechanistic framework for understanding how rapid ecological differentiation can occur under strong genomic constraint and may represent a general evolutionary strategy in complex eukaryotic microbes.

## Results

### A conserved genomic backbone underlies ecological divergence

Phylogenomic analysis using 8,653 single-copy orthologs resolved eight *Metarhizium robertsii* strains into two well-supported clades corresponding to early- and recently diverged lineages (Fig. 1). This phylogenetic structure aligned with previously reported phenotypic differentiation (Sheng and St. Leger 2026), providing a framework to investigate genomic changes accompanying ecological differentiation.

**Fig. 1.**
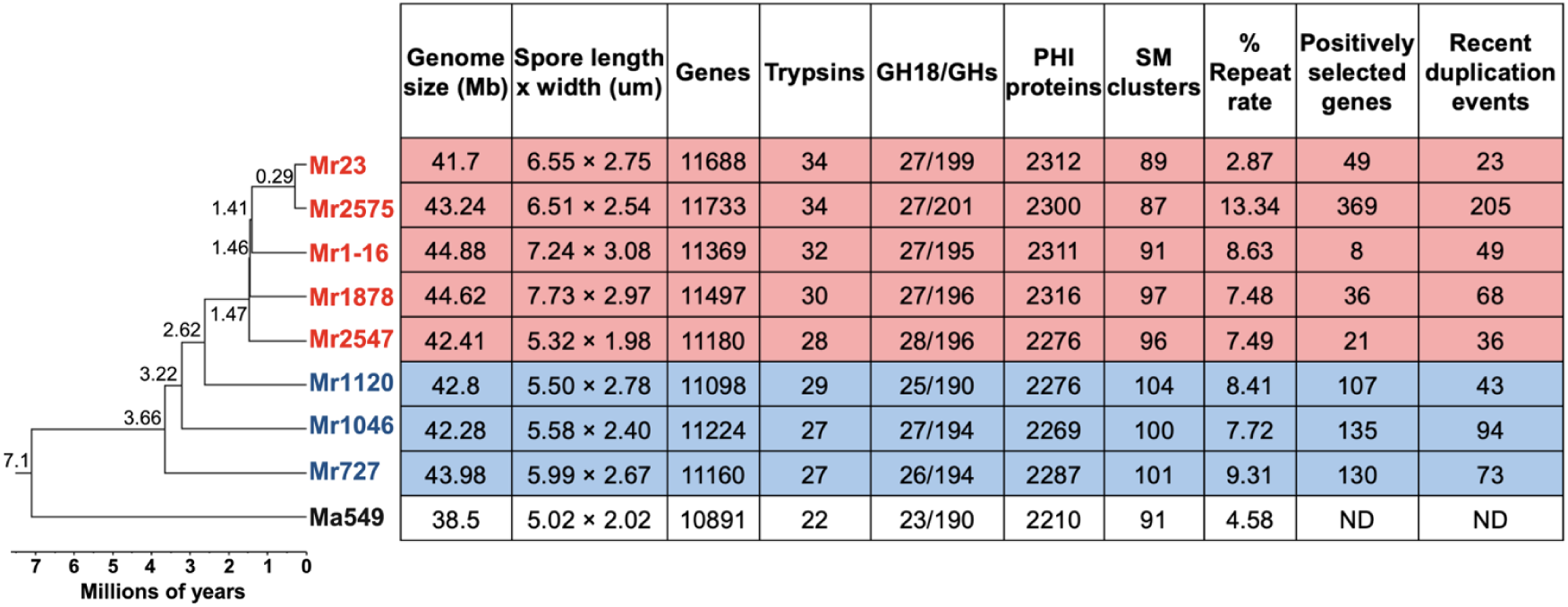
Phylogenomic relationships and genome feature variation among eight *M. robertsii* strains. Maximum-likelihood phylogeny inferred from 8,653 single-copy orthologs, with *M. anisopliae* ARSEF549 (Ma549) as the outgroup (Hu et al. 2014). The strains resolve into two clades corresponding to early diverged (shown in blue; Mr727, Mr1046, Mr1120) and recently diverged (shown in red; Mr1878, Mr2547, Mr1-16, Mr2575, Mr23) lineages. Genome size, total gene number, and spore dimensions are shown adjacent to the tree. Functional categories associated with strain diversification are summarized, including trypsins, GH18 chitinases, pathogen–host interaction (PHI) genes, and secondary metabolite (SM) gene clusters. Genome repeat content, numbers of positively selected genes, and recently duplicated events unique to individual strains are also indicated. ND, not determined.

Despite clear ecological divergence, genome architecture was remarkably conserved. Average nucleotide identity (ANI) exceeded 98.5% among all strains (Table S1), and whole-genome alignments revealed extensive macrosynteny (Fig. 2; Fig. S1). Gene order conservation extended beyond the species level, with substantial synteny (96.5%) retained relative to its sister taxon *M. anisopliae* ARSEF549 (Ma549), which diverged approximately 7.1 MYA (Fig. 1). Notably, synteny between *M. robertsii* and *Pochonia chlamydosporia* remained high (79.88%) despite ∼180 MY of divergence, indicating long-term preservation of gene order within this lineage, a pattern uncommon in fungi(Hane et al. 2011).

**Fig. 2.**
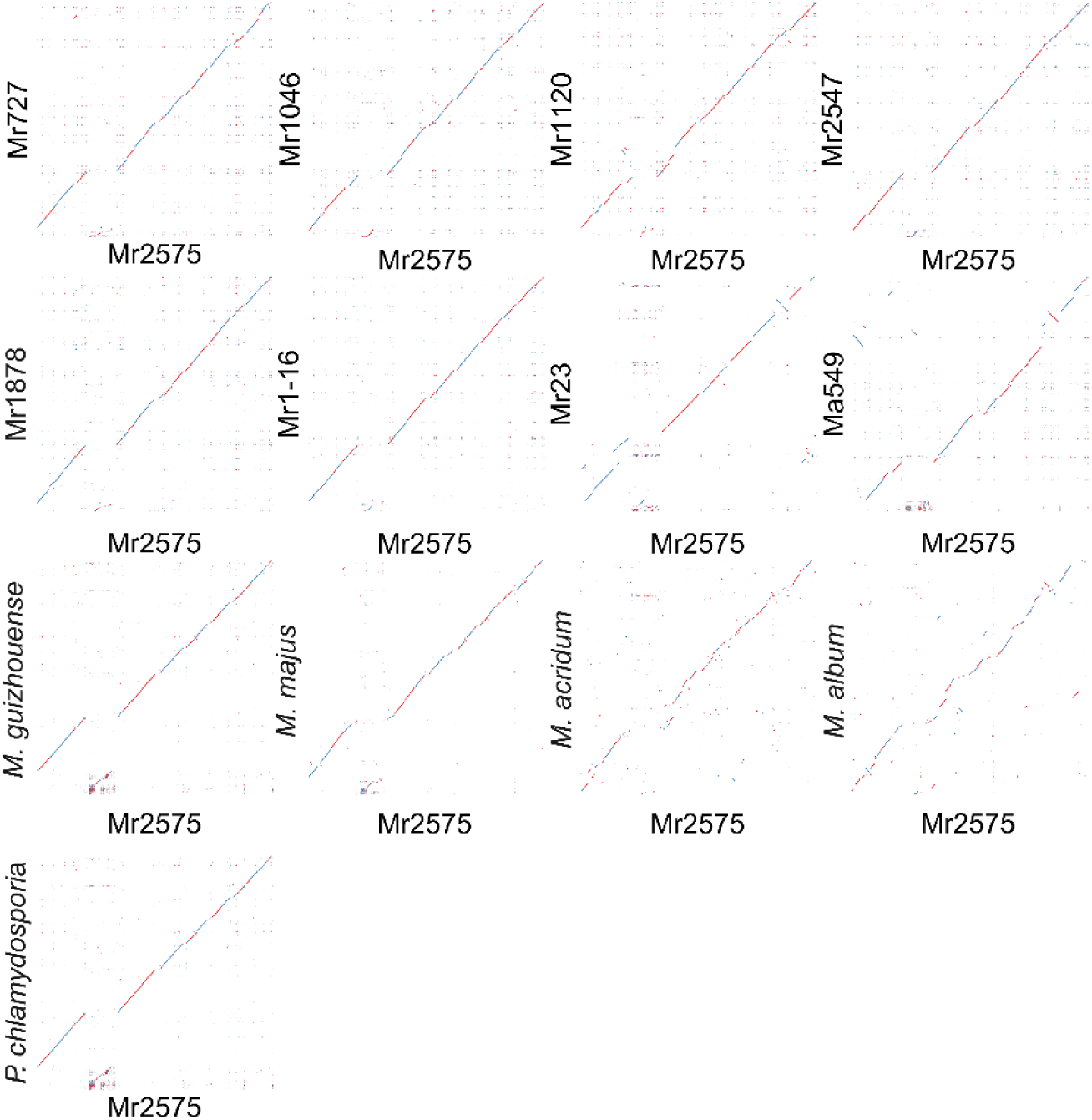
Whole-genome synteny comparisons between Mr2575 and other *M. robertsii* strains and related fungi. Dot plots show pairwise genome alignments generated using NUCmer in MUMmer v4.0.0(Marçais et al. 2018). Alignments are shown between Mr2575 and other *M. robertsii* strains, additional *Metarhizium* species, and *P. chlamydosporia*. Scaffolds from Mr2575 assembly are plotted on the x-axis, and scaffolds from the comparison genome are shown on the y-axis. Diagonal alignments indicate regions of sequence similarity; blue lines represent forward-strand alignments and red lines represent reverse-strand alignments. Whole-genome synteny comparisons using Mr23 as the reference genome are shown in Fig. S1.

All eight *M. robertsii* assemblies were highly complete (>97%; Table S2) ensuring that observed differences reflect biological variation rather than technical artifacts. Compared with early diverged specialist species (*M. album* and *M. acridum*), *M. robertsii* strains had significantly larger genomes (43.24 Mb vs. 34.95 Mb; p = 0.0029) and encoded more protein-coding genes (11,369 vs. 9,161; p = 5.067e-5) (Table 1). Among *M. robertsii* strains, recently diverged lineages encoded slightly more genes than early diverged strains (11,493 vs. 11,161; Welch’s t test, p = 0.0270) (Fig. 1 and Table 1) (Table S3). Approximately 88.6% of genes are shared across all genomes (range: 85.6–89.8%), with the remaining genes constituting a non-core fraction, comprising accessory genes present in subsets of strains and a small number (9–73 per strain) of strain-specific genes (Table S3). Notably, more than 58% of the strain specific genes were single-copy orphans, potentially therefore arising through gene duplication, horizontal gene transfer (HGT), or rapid sequence divergence (McCarthy and Fitzpatrick 2019).

**Table 1.**
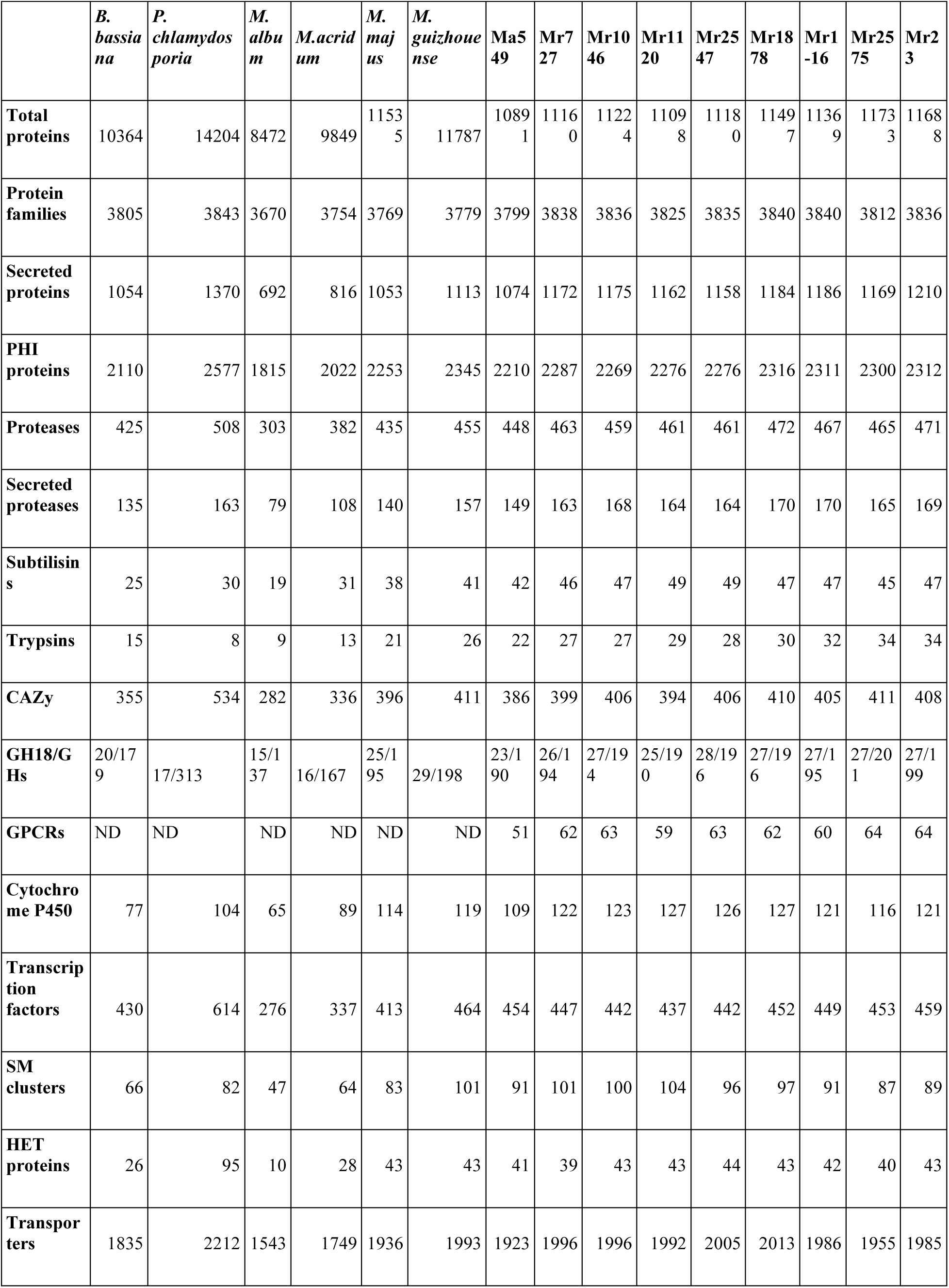
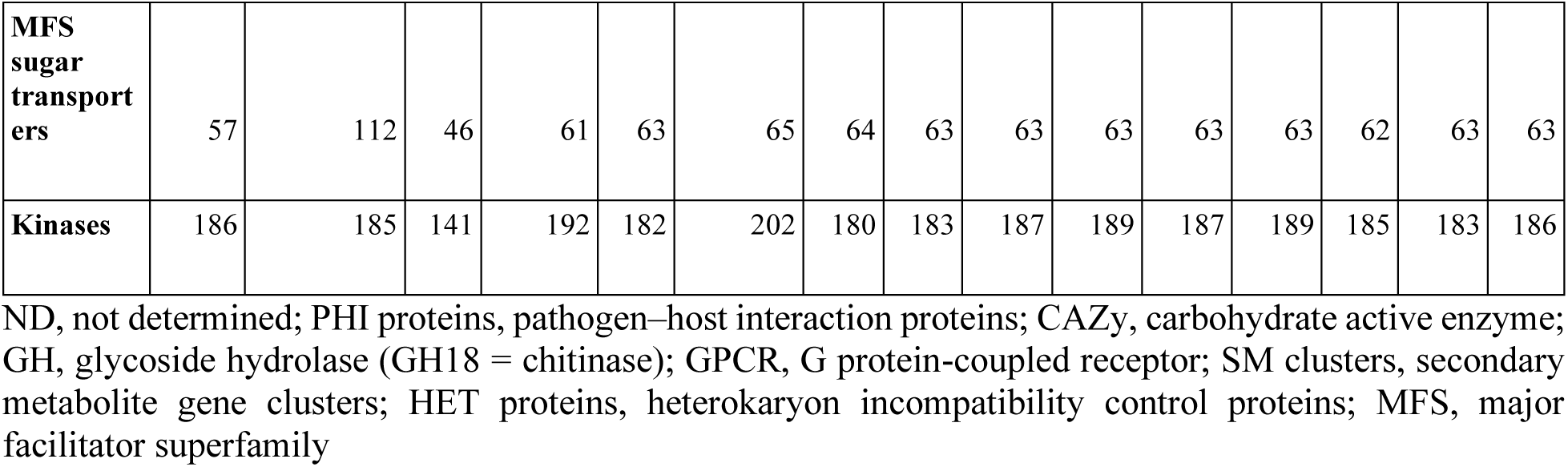
Comparative genome features and protein family composition across *Metarhizium* and related fungi.

Phylogenetically independent contrast (PIC) analysis of *M. robertsii* strains revealed significant correlations between genome size and gene-coding capacity (F = 8.20, P = 0.0174), as well as between genome size and spore morphology (length: F = 23.292, P = 0.0029; width: F = 18.43, P = 0.0051). Gene number was likewise significantly correlated with spore dimensions (length: F = 30.158, P = 0.0015; width: F = 13.628, P = 0.01). These patterns suggest coordinated evolution of genome expansion and spore morphological traits.

### Genomic divergence involves repeat-rich and dynamic regions

Two accessory chromosomes (chrA and chrB) have been described in *M. robertsii* as horizontally transferable genomic elements that contribute to fitness and adaptation(Habig et al. 2024). However, non-overlapping alignment coverage against these accessory chromosome references indicated that neither chrA nor chrB is maintained as an intact chromosome across *M. robertsii* strains but are instead fragmented and redistributed across the genomes.

In contrast to the conserved core genome architecture, genomic divergence was strongly localized to repeat-rich regions. We intersected synteny gap intervals with sequences homologous to chrA and chrB. Across pairwise comparisons involving Mr2575, 17.4–32.4% of gap bases overlapped accessory-derived regions (Table S4), indicating that while these sequences contribute to localized divergence, the majority (∼70–80%) occurs outside accessory chromosome remnants.

Synteny gaps between closely related strains were highly enriched for repetitive elements; for example, in the comparison between Mr23 and Mr2575, 68.5% of gap sequences overlapped repeats (Fig. 2; Fig. S1). These discontinuities were not randomly distributed but instead occurred at shared genomic loci across independent comparisons. Using Mr2575 as a reference coordinate system, we quantified the extent to which gap regions identified in one pairwise comparison were shared with those identified in another (Table S5). The proportion of shared gap loci ranged from 25.1% to 84.0%. Across all comparisons, the extent of sharing exhibited a weak negative association with sequence divergence (Spearman’s ρ = −0.265, P = 0.245), although this relationship was not statistically significant. Together, these results indicate that genomic divergence involves a restricted set of repeat-rich regions that are recurrently modified across strains, though the spatial distribution of repeat accumulation varies substantially among lineages.

To assess whether these divergence-associated regions were enriched near chromosome ends, we intersected synteny gap intervals with subtelomeric regions defined by proximity to telomeric repeat tracts. Across pairwise comparisons, subtelomeric regions comprised approximately 16–21% of the genome but accounted for ∼30–50% of synteny gap bases, indicating significant enrichment of divergence near chromosome ends (Table S6). However, this enrichment was not uniform across all strains and was influenced by assembly contiguity. More fragmented assemblies exhibited inflated subtelomeric coverage due to increased numbers of scaffold ends, whereas more contiguous assemblies (e.g., Mr23 and Mr2575) showed reduced subtelomeric fractions despite comparable or higher telomere detection rates. These patterns indicate that variation in subtelomeric coverage primarily reflects assembly structure rather than biological differences. These results indicate that while chromosome-end–proximal regions act as recurrent hotspots of structural variation, divergence is not confined to subtelomeric compartments but is distributed across shared, repeat-associated loci throughout the genome.

This spatially restricted pattern of variation explains how extensive macrosynteny is maintained despite fragmentation and redistribution of accessory chromosome–derived sequences. Coverage of chrA ranged from 15.6% (Mr1120) to 67.3% (Mr2575), with chrB showing a similar range (15.9% in Mr1120 to 57.7% in Mr2575; Table S7). In Mr2575, which retained among the highest proportions of both chromosomes, homologous sequences were distributed across multiple PacBio long-read contigs (Fig. S2). For chrA, alignments were concentrated in contigs 10 and 13, mapping primarily to ∼0.45–0.69 Mb and ∼0.76–1.22 Mb, respectively, as multiple discrete, locally collinear blocks separated by gaps. Similarly, chrB-derived sequences were distributed across contigs 12, 14, and 16, forming clusters of locally collinear blocks interspersed with shorter more dispersed alignments. While chrA and chrB are enriched in transposable elements (30.96% and 32.48%, respectively) and exhibit low gene densities (∼22%)(Habig et al. 2024), the corresponding contigs in Mr2575 showed lower TE content (6.47–18.54%) and higher gene densities (25.49–30.01%; Table S8), indicating substantial restructuring of accessory chromosome-derived sequences.

Variation in repetitive content was dominated by transposable elements (TEs) (Table S9). TE content varied markedly among strains despite minimal nucleotide divergence, with even closely related strains exhibiting substantial differences in TE loads and element length. Thus, TE abundance ranged from less than 3% in Mr23 to over 13% in Mr2575, despite these strains diverging only ∼0.29 MYA (Table S9). These dynamics translate to remarkably rapid evolutionary rates: TE proliferation in Mr2575 reached ∼43 elements per million years (assuming 0.29 MY divergence), while RIP-mediated genome defense varied from ∼0.9% genome affected per MY in Mr23 to ∼13.8% per MY in Mr1120. The 10-fold variation in TE loads across such shallow phylogenetic distances (from <3% to >13% total genomic content) represents one of the most rapid documented cases of repeat expansion within fungal species complexes, highlighting the potential for transposable elements to drive genome evolution over ecological timescales rather than requiring deep evolutionary time.

TEs in Mr2575 were also significantly longer (mean ± SE: 409 ± 5 bp) than those in closely related strains Mr1-16 (292 ± 4 bp) and Mr23 (180 ± 4 bp) (Kruskal–Wallis H = 1281.8, p < 2.2e-16) (Table S10). Since TE fragments typically shorten over time due to mutation and deletion (Maumus and Quesneville 2014), the enrichment of longer elements in Mr2575 is consistent with relatively recent transposition activity. Notably, the elevated TE activity in Mr2575 was not associated with increased subtelomeric sequence content, which comprised only ∼5% of the genome similar to Mr23, in contrast to the ∼20% observed in other strains sequenced with Illumina alone. This distribution suggests that TE proliferation in Mr2575 occurred predominantly in internal chromosomal regions rather than subtelomeric compartments, indicating strain-specific variation in the spatial dynamics of repeat expansion and potentially different regulatory constraints on TE insertion sites across the genome. Family-level comparisons further showed expansion of multiple TE families in Mr2575 and contraction in Mr23 relative to Mr1-16 (Fig. 3). Phylogenetic analyses indicated that these expansions primarily involved proliferation of resident TE families rather than acquisition of novel elements, consistent with lineage-specific activation of endogenous repeats rather than horizontal acquisition (Figs. S3–S4).

**Fig. 3.**
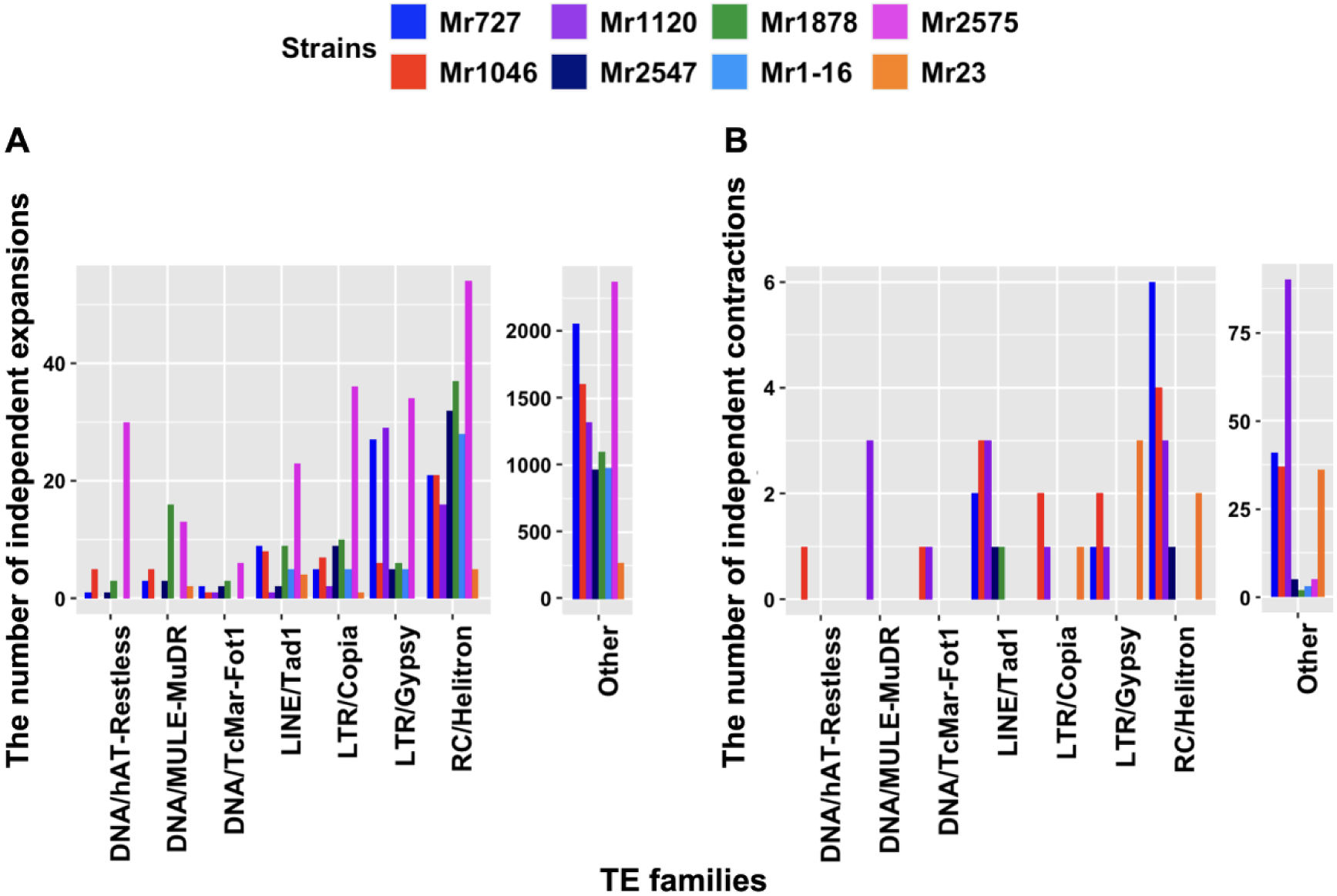
Transposable element abundance variation across *M. robertsii* genomes. LTRs, Long Terminal Repeats; LINEs, Long Interspersed Nuclear Elements; SINEs, Short Interspersed Nuclear Elements.

### Repeat-Induced Point Mutation Contributes to Genomic Divergence

Variation in repeat-induced point mutation (RIP), a genome defense mechanism that targets duplicated DNA(Lewis et al. 2010; Gessaman and Selker 2017; He et al. 2020), further contributed to genomic divergence. Although the core RIP machinery was conserved across all strains (Fig. 4; Table S11), the extent of RIP activity varied substantially. Using both RIPCAL (Fig. S5) and the stricter RIPper criteria (RIP product > 1.15; RIP substrate ≤ 0.75; RIP composite index > 0), we quantified RIP-affected windows and large RIP-affected regions (LRARs) following established protocols (Van Wyk et al. 2019; Van Wyk et al. 2021).

**Fig. 4.**
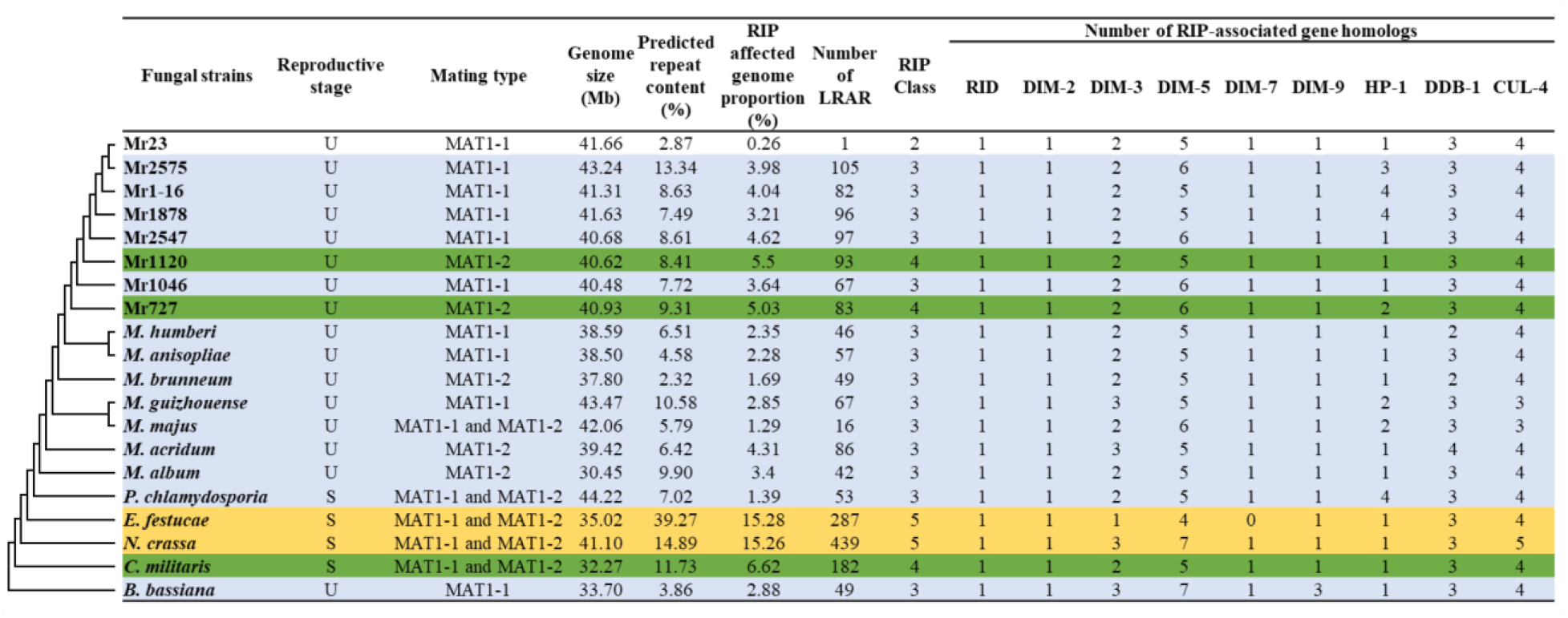
Comparative analysis of repeat-induced point mutation (RIP) across *M. robertsii* and related fungi. Left, cladogram showing phylogenetic relationships among the analyzed fungi, with *Beauveria bassiana* as the outgroup. Adjacent columns summarize reproductive strategy (S, sexual; U, unknown), mating type, genome size (Mb), percentage of repetitive DNA, total genomic fraction affected by RIP, and number of large RIP-affected regions (LRARs). RIP intensity classes are indicated by color shading (Class 2: 0.2–1.0%; Class 3: 1.0–5.0%; Class 4: 5.0–10.0%; Class 5: 10.0–20%)(Van Wyk et al. 2021). Right, presence of homologs to canonical RIP-associated genes, including RID, DIM-2, and associated cofactors (HP-1, DIM-3, DIM-5, DIM-7, DIM-9, DDB-1, and CUL-4) characterized in *Neurospora crassa* (Lewis et al. 2010; Gessaman and Selker 2017; He et al. 2020).

Early diverged strains Mr1120 and Mr727 exhibited higher fractions of RIP-affected genome (5.5% and 5.03%, respectively) and grouped with sexually reproducing comparators such as *Cordyceps militaris* (Fig. 4)(Class 4 as categorized by Van Wyk et al. 2021), consistent with retention of RIP activity or recent sexual history (Van Wyk et al. 2021). In contrast, Mr23 showed minimal RIP signal (0.26% RIP-affected genome), while remaining strains showed moderate RIP (1–5% RIP-affected genome) (Fig. 4; Table S12). Across the broader taxon set TE abundance, RIP signatures, and repeat distribution were strongly correlated (Spearman r = 0.75–0.87, p ≤ 0.0002) (Fig. 4; Table S12), indicating tight coupling between transposable element activity and genome defense processes.

TE classes differed markedly in their susceptibility to RIP. LINE and Helitron elements were rarely detected within RIP-affected regions, whereas LTR retrotransposons (Gypsy and Copia families) were disproportionately enriched (Fig. 5; Table S13), consistent with preferential targeting of specific repeat types. Among strains, the proportion of TEs impacted by RIP varied widely from high values in Mr1120 (47.67% of TEs in RIP-affected windows; 28.5% in LRARs) to very low values in Mr23 (2.51% in RIP-affected windows; 0% in LRARs), mirroring their overall TE loads and inferred RIP histories. Notably, Mr2575 combined the highest TE burden (12.49% of the genome) with only moderate RIP activity (3.98% RIP affected genome), consistent with incomplete TE suppression rather than strong genome defense. This suggests that the balance between TE proliferation and genome defense mechanisms varies among strains and may contribute to lineage-specific evolutionary trajectories.

**Fig. 5.**
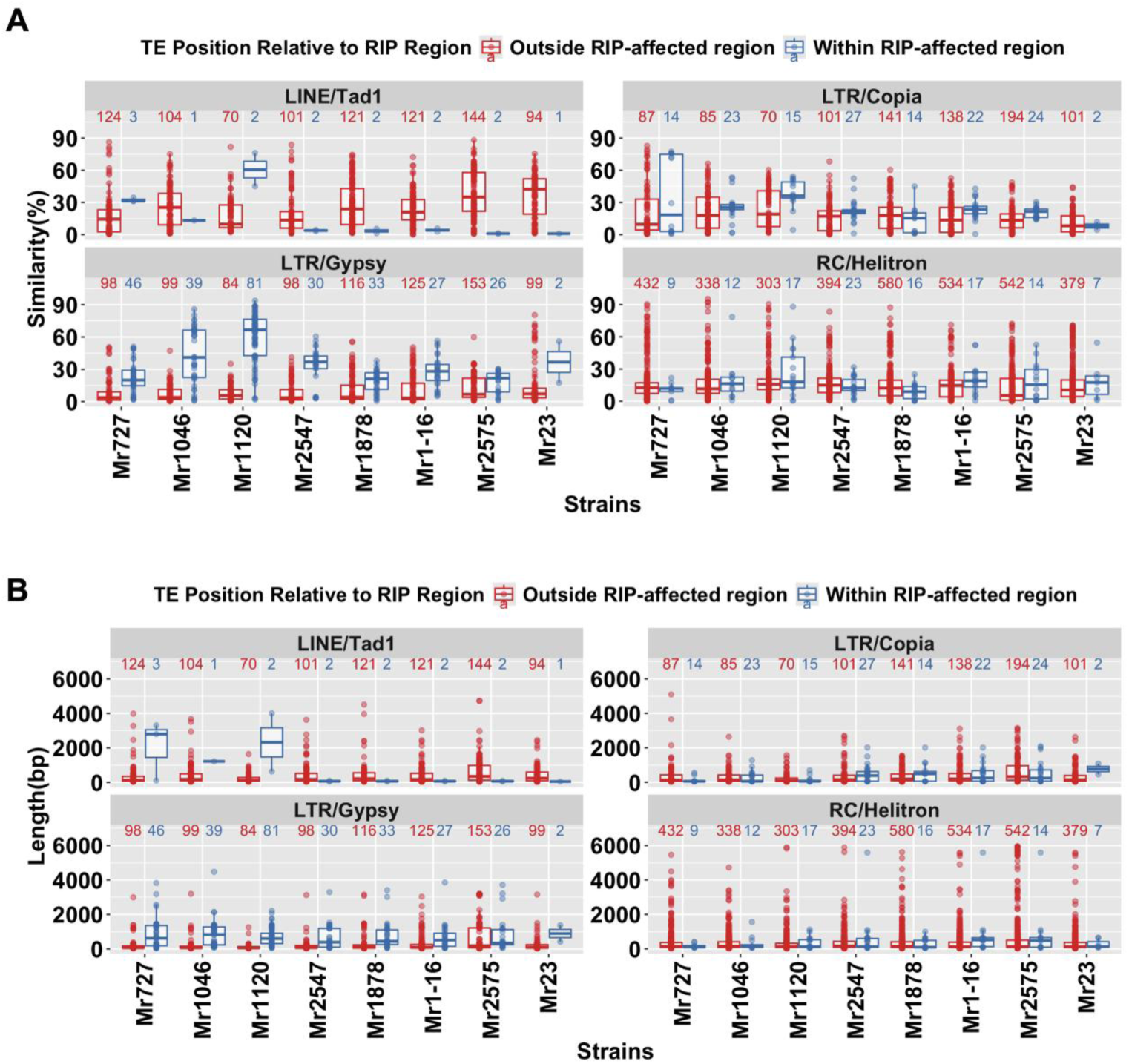
Transposable element similarity and length distributions in RIP-affected and non-RIP genomic regions. Similarities were calculated based on consensus sequence of each TE family in each *M. robertsii*. The number of TEs in each family in RIP-affected and non-RIP regions were indicated on top of each box.

### Mobile DNA dynamics couples with gene duplication and positive selection

Patterns of gene duplication closely parallel variation in TE content (Table S2). *M. robertsii* strains with higher TE abundance exhibited significantly greater numbers of recently duplicated genes, with a strong positive correlation between TE load and duplication frequency (r = 0.79, p = 0.02). Across all strains, duplicated genes were significantly enriched within 5 kb of TEs compared with the genomic background (Table S14, 39–68% vs. 10–14%; Fisher’s exact test, FDR < 1e-11). These results support a model in which TE-rich genomic regions act as hotspots for recent gene duplication. Lineage-specific gene duplications and numbers of positively selected genes (PSGs) were unevenly distributed across strains, with recently diverged lineages showing greater heterogeneity than earlydiverged strains (Table 2). PSG counts strongly correlated with duplication events (*r* = 0.93, *P* = 0.0007) indicating that both processes contribute to lineage-specific adaptation. Mr2575 exhibited the highest numbers of TEs, duplicated genes, and PSGs, whereas Mr23 showed the lowest TE content and duplication counts, and Mr1-16 harbored the fewest PSGs. These patterns indicate that both gene duplication and positive selection have become increasingly heterogeneous in recently diverged lineages, consistent with divergence in selective pressures among strains. Because PSGs were identified only from orthologs shared across all strains, i.e., genes duplicated prior to strain divergence, recently duplicated genes unique to individual strains were excluded from PSG analyses. Accordingly, gene duplication and positive selection represent coordinated but largely non-overlapping processes contributing to lineage-specific adaptation.

**Table 2.** Positively selected genes and genes derived from recent lineage-specific duplication events across functional categories, including PHI-base orthologs in parenthesis.

|  | Protein families (PHI) | Mr727 | Mr1046 | Mr1120 | Mr2547 | Mr1878 | Mr1-16 | Mr2575 | Mr23 |
| --- | --- | --- | --- | --- | --- | --- | --- | --- | --- |
| **CAZymes** | **PSG** | 4 (3) | 9 (4) | 4 (2) | 0 | 0 | 1 (1) | 10 (2) | 1 (0) |
|  | **duplicated genes** | 3 (2) | 5 (2) | 0 | 0 | 2 (2) | 1 | 17 (6) | 0 |
| **Proteases** | **PSG** | 13 (1) | 12 (3) | 4 (1) | 1 (0) | 4 (2) | 0 | 15 (3) | 3 (0) |
|  | **duplicated genes** | 7 (2) | 2 (0) | 3 (0) | 2 (0) | 3 (0) | 1 (0) | 3 (0) | 0 |
| **Total Transporters** | **PSG** | 46 (16) | 39 (10) | 29 (11) | 9 (5) | 17 (8) | 3 (3) | 126 (58) | 19 (10) |
|  | **duplicated genes** | 23 (4) | 26 (7) | 20 (9) | 6 (1) | 7 (0) | 2 (0) | 49 (15) | 7 (3) |
| **ABC transporters** | **PSG** | 0 | 0 | 0 | 1 | 0 | 0 | 3 | 0 |
|  | **duplicated genes** | 4 (2) | 5 (3) | 0 | 1 | 0 | 0 | 0 | 0 |
| **MFS transporters** | **PSG** | 5 (1) | 9 (2) | 3 (0) | 0 | 1 (0) | 1 (1) | 12 (6) | 4 (2) |
|  | **duplicated genes** | 0 | 2 (1) | 6 (2) | 0 | 0 | 0 | 5 (1) | 0 |
| **Secondary metabolite core genes** | **PSG** | 7 (3) | 6 (3) | 3 (1) | 0 | 2 (1) | 0 | 5 (1) | 4 (1) |
|  | **duplicated genes** | 5 (2) | 2 (1) | 3 (2) | 1 | 2 | 0 | 0 | 1 (1) |
| **Cytochrome P450\*** | **PSG** | 4 (3) | 4 (1) | 1 (1) | 0 | 1 (1) | 0 | 2 (2) | 2 (1) |
|  | **duplicated genes** | 0 | 4 (1) | 7 (5) | 2 (1) | 0 | 0 | 0 | 2 (2) |
| **Transcription factors** | **PSG** | 7 (1) | 5 (1) | 5 (0) | 1 (0) | 1 (0) | 1 (0) | 17 (5) | 1 (0) |
|  | **duplicated genes** | 4 (1) | 6 (1) | 2 (0) | 1 (0) | 0 | 0 | 8 (2) | 1 (1) |
| **Sum** | **PSG** | 130 (27) | 135 (31) | 107 (23) | 21 (7) | 36 (17) | 8 (3) | 369 (114) | 49 (15) |
|  | **duplicated genes** | 139 (24) | 177 (13) | 84 (10) | 68 (1) | 127 (8) | 88 (4) | 368 (37) | 49 (5) |

\*Six out of 27 positively selected secondary metabolites core genes are also characterized as positively selected cytochrome P450s.

### Positive selection predominately targets intracellular and regulatory functions

Positive selection and gene duplication events were disproportionally associated with intracellular processes, except in Mr727, where secreted genes were significantly more likely to be duplicated (23/139; Fisher’s exact test, P = 0.025, odds ratio = 1.70). Overall, only about 8% of PSGs encoded secreted proteins (Table 2), indicating that extracellular effector repertoires are not the primary targets of divergence (Fig. S6).

Approximately 70% of PSGs assigned to biological processes were strain-specific (Tables S15–S16), consistent with lineage-specific adaptation. Nevertheless, several strains showed enrichment for similar functional categories, suggesting repeated targeting of related pathways. Transport-related functions were a recurrent target of selection. PSGs in Mr23, Mr2575, Mr1878, and Mr727 were significantly enriched for transporter-related functions (Fisher’s exact test, p < 0.05), consistent with adaptation involving substrate uptake, efflux, or homeostasis. In Mr2575, 126 of 369 PSGs were distributed across 70 transporter families (Tables S17-18), typically represented by single genes, indicating dispersed rather than cluster-based expansion. Comparative analysis of the PF00083 PFAM (“Sugar and other transporters”) as a representative large family further highlighted ecological specialization. The related species *Trichoderma reesei*, known for plant cell wall degrading enzymes, encodes a broader repertoire of plant cell wall–associated transporters, whereas *M. robertsii* strains possess additional trehalose-associated transporters, consistent with adaptation to trehalose as the principal insect hemolymph sugar (Fig. S7).

Multiple strains (Mr1878, Mr23, and Mr727) also showed functional enrichments in signaling-related activities (GTPase, protein kinases, regulator domains) consistent with diversification of environmental sensing. Different strains also showed distinct functional enrichments (Tables S15-16). Mr2575 PSGs were enriched for RNA binding and structural molecule activity, including RNA recognition motifs, implicating adaptive evolution in transcriptional regulation, RNA processing, and translation. Mr727 and Mr2575 shared enrichment for protein maturation indicating selection on post-translational processing. Mr1046 PSGs were enriched for oxidoreductase activity, including cytochrome P450s, suggesting selection on detoxification and oxidative metabolism. Mr2547 PSGs were enriched for lyase activity, implicating carbon metabolism and detoxification pathways. Only Mr727 and Mr1046 showed enrichment for chromosome organization, including topoisomerases, indicating lineage-specific selection on genome stability.

Analysis of PSG and lineage-specific duplicated genes with homology to entries in the Pathogen–Host Interaction (PHI) database reinforced these patterns. Approximately 20.2% of *M. roberstii* genes were PHI-associated and these were broadly conserved across strains (Table 1), with approximately 75% classified in the “loss of pathogenicity” and “reduced virulence” PHI categories (Table S19). On average, 30.6% of PSGs and 8.6% of duplicated genes were PHI-associated, indicating that adaptive evolution of host interaction traits is driven more strongly by positive selection than by duplication (Table S20).

Despite this, few well characterized *M. robertsii* virulence genes showed evidence of selection. Canonical factors, including the adhesins Mad1 and Mad2 (Wang and St. Leger 2007), and the collagen-like immune evasion protein MCL1 (Wang and St. Leger 2006), were highly conserved and lacked signatures of positive selection. Similarly, fatty acid–binding proteins (FABP1 and FABP2) and sugar transporters (Mst1 and Mrt) implicated in rhizosphere and rhizoplane colonization (Dai et al. 2021), were conserved across all strains without evidence of selection. In contrast, five PSGs (one in Mr1046, three in Mr2575, and one in Mr23) exhibited moderate homology to Mst1 (16–27%) or Mrt (14–20%) but more closely resembled STL1-like glycerol transporters, inositol transporters, and maltose permeases. Given that STL1 mediates glycerol accumulation under osmotic stress in other fungi (Ferreira et al. 2005), these signals of positive selection likely reflect adaptation to osmotic or host-associated environments.

PSGs in Mr1046 and Mr727 were enriched for protease-associated functions, with a substantial fraction predicted to be secreted (5/13 and 7/12, respectively), consistent with strain-specific optimization of host interaction or nutrient acquisition. Only two GH18 chitinases showed signatures of positive selection (in Mr1046 and Mr727), one of which (Mr727) is predicted to be secreted (Fig. S8). In contrast PSGs in Mr23 were enriched for secondary metabolism core genes, indicating adaptive diversification of metabolic pathways. In Mr2575 and Mr727, duplication generated additional small cysteine-free proteins (CFPs), a class associated with virulence in Mr23 (Mou et al. 2022).

### Gene content innovation and horizontal gene transfer

Horizontal gene transfer (HGT) appears to have contributed only modestly to intraspecific diversification in *M. robertsii*. Most previously identified HGT-derived genes in Mr23 (Zhang et al. 2019) were conserved across all strains, indicating that these acquisitions likely predate strain divergence. Nonetheless, a small number of strain-restricted genes suggest that more recent, lineage-specific acquisitions have occurred. These include a defensin (PF01097) in Mr727, similar (e-value 1e-18; 57% identity) to the anti-bacterial scedosporisin-2 from a wide-spread soil fungus (Wu et al. 2018), consistent with occasional contributions of external genetic material linked to ecological adaptation.

CYP450 gene families are notably expanded in *M. robertsii* (average: 123) compared to other fungi (Table 1), consistent with the metabolic versatility required for its multifunctional lifestyle. Several CYP genes may have been horizontally acquired (Table S21). Most *Metarhizium* species, including *M. robertsii,* have two CYP55 genes, whereas *P. chlamydosporia, M. album* and *B. bassiana*, like most fungi (Chen et al. 2014), lack these genes. CYP55 enzymes catalyze the reduction of nitric oxide (NO) to nitrous oxide (N_₂_O) which facilitates survival in hypoxic or anoxic environments, potentially relevant to both insect cadavers and soil niches.

Additional CYP lineages show patchy phylogenetic distributions suggestive of complex evolutionary histories. A putative CYP105 gene is present as a single copy in most *M. robertsii* strains but absent from Mr1046 and Mr727. Its closest homologues occur in the insect pathogen *Torrubiella hemipterigena* (79%) and the metal-tolerant ericoid mycorrhizal fungus *Oidiodendron maius* (78%), whereas similarity to functionally characterized proteins is low (∼31% identity to a linoleate diol synthase from *Magnaporthe grisea*(Cristea et al. 2003). This pattern is consistent with either ancient acquisition followed by divergence or differential retention across lineages and does not by itself resolve origin.

By contrast, CYP57 shows a more restricted distribution, being present only in Mr727, Mr1878 and *M. majus* (>99% similarity). This protein shares moderate similarity (37%) with a *Fusarium oxysporum* pisatin demethylase involved in detoxification of the phytoalexin pisatin (Maloney and VanEtten 1994) (Table S21). Given that this detoxification gene family has been inferred to spread within *Fusarium* via HGT (Milani et al. 2012), the distribution observed here is compatible with a scenario in which a related gene was acquired prior to the divergence of *M. majus* and *M. robertsii*, followed by lineage-specific loss.

### Core functional repertoires are conserved despite ecological differentiation

Despite pronounced ecological divergence, major functional gene classes remained highly conserved across *M. robertsii* strains, supporting the localized innovation model.

### Limited protease divergence despite virulence differences

Proteases represent one of the most expanded functional classes in *Metarhizium*, consistent with insect-derived nutrition (St. Leger and Wang 2020). Relative to early diverged specialists, such as *M. acridum* and *M. album*, *M. robertsii* encodes substantially larger protease repertoires (mean 471 vs. 342), including expansions of serine, aspartic, and metalloproteases (Tables S22–S23) (St. Leger et al. 1987; Bagga et al. 2004; Hu et al. 2014). However, despite marked ecological divergence among strains, these major functional classes showed minimal variation in both composition and size across strains, indicating that expansion of these families predates intraspecific diversification.

Limited gene turnover occurred within larger protease families. For example, trypsins exhibited balanced dynamics across the eight *M. robertsii* strains (ten gains and ten losses; (Figs. S9 and S10), consistent with lineage-specific duplication and loss rather than directional expansion. However, a different pattern is observed for metalloprotease M16A (pitrilysin), which functions in a-factor mating pheromone processing (Alper et al. 2006). While most strains encode a single copy, Mr1878 and Mr727 contain five and four copies, respectively, reflecting recent duplication in these lineages coupled with gene loss in others (Fig. S11).

At finer phylogenetic resolution, several MEROPS families display strain-restricted distributions. The Kazal-type serine protease inhibitor family is detected only in Mr2575 and Mr23, whereas family I51, implicated in regulatory signaling processes, is confined to Mr1120 and Mr727. Likewise, the cysteine peptidase family C45, that is common in fungi and associated with secondary metabolism, is present only in Mr23 and Mr2575 (Table S22). Together, these patterns indicate that while the overall protease arsenal is evolutionarily stable, localized duplication and loss contribute to functional diversification.

### Metabolic specialization through functional repurposing

Carbohydrate-active enzyme (CAZyme) repertoires were similarly stable across *M. robertsii* strains. Although *M. robertsii* lacks several canonical cellulase and xylanase families characteristic of plant biomass decomposers such as *P. chlamydosporia* and *Metarhizium marquandii* (Tables 1, 3 and 4; Table S24), it retains a consistent complement of enzymes targeting pectin and other non-cellulosic substrates (Table 3; Table S24). Differences between early and recently diverged strains were minimal, indicating that strain-level ecological divergence between plant associates and non-plant associates(Sheng and St. Leger 2026) is not driven by large-scale remodeling of degradative enzyme systems (Tables 1, 3 and 4; Table S24). Notable strain-to-strain differences were limited to specific oxidative or chitin-associated families, including modest differences in chitin active AA11 lytic polysaccharide monooxygenases and GH75 chitosanase counts, rather than systematic expansion of plant cell wall hydrolases.

**Table 3.**
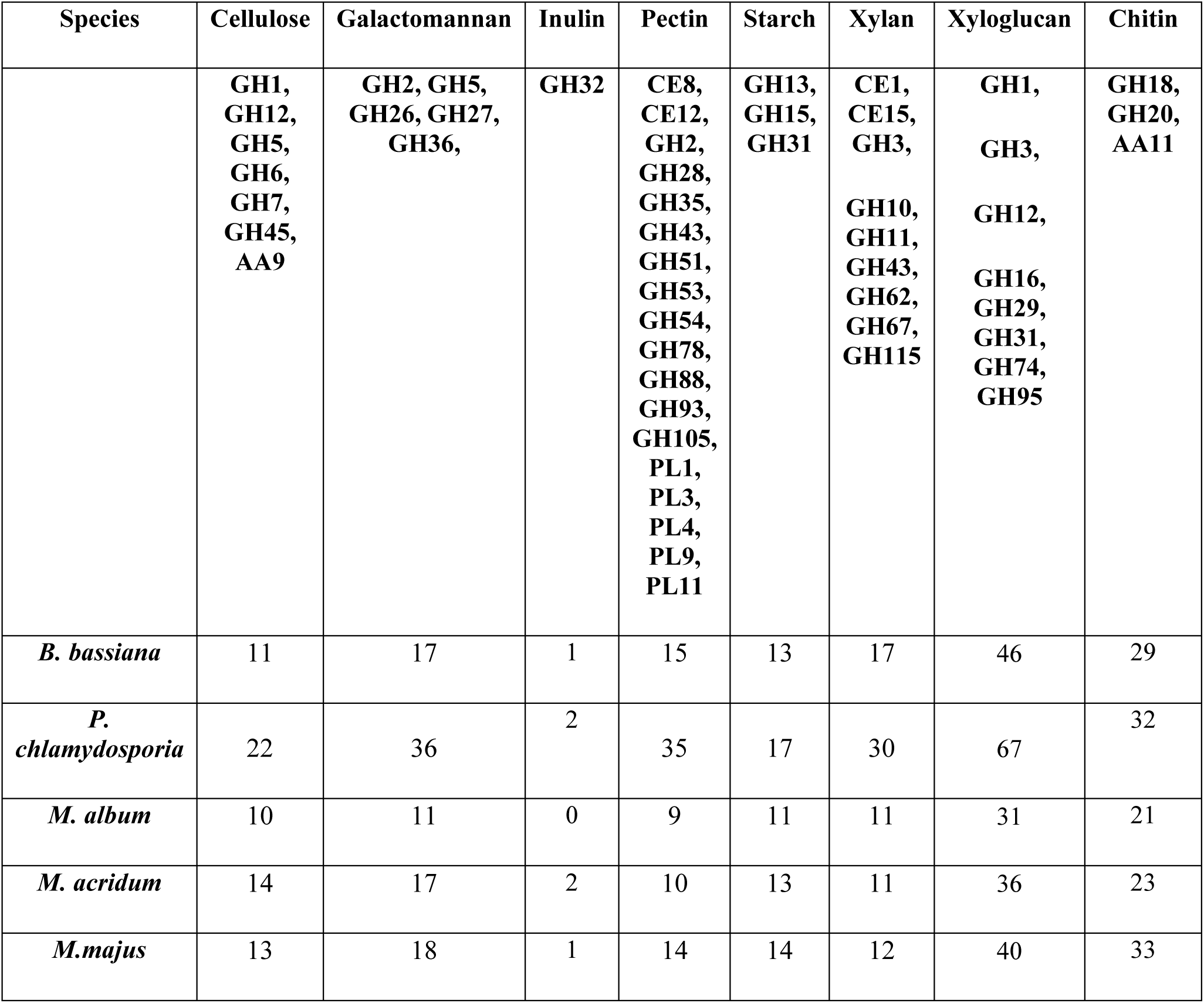
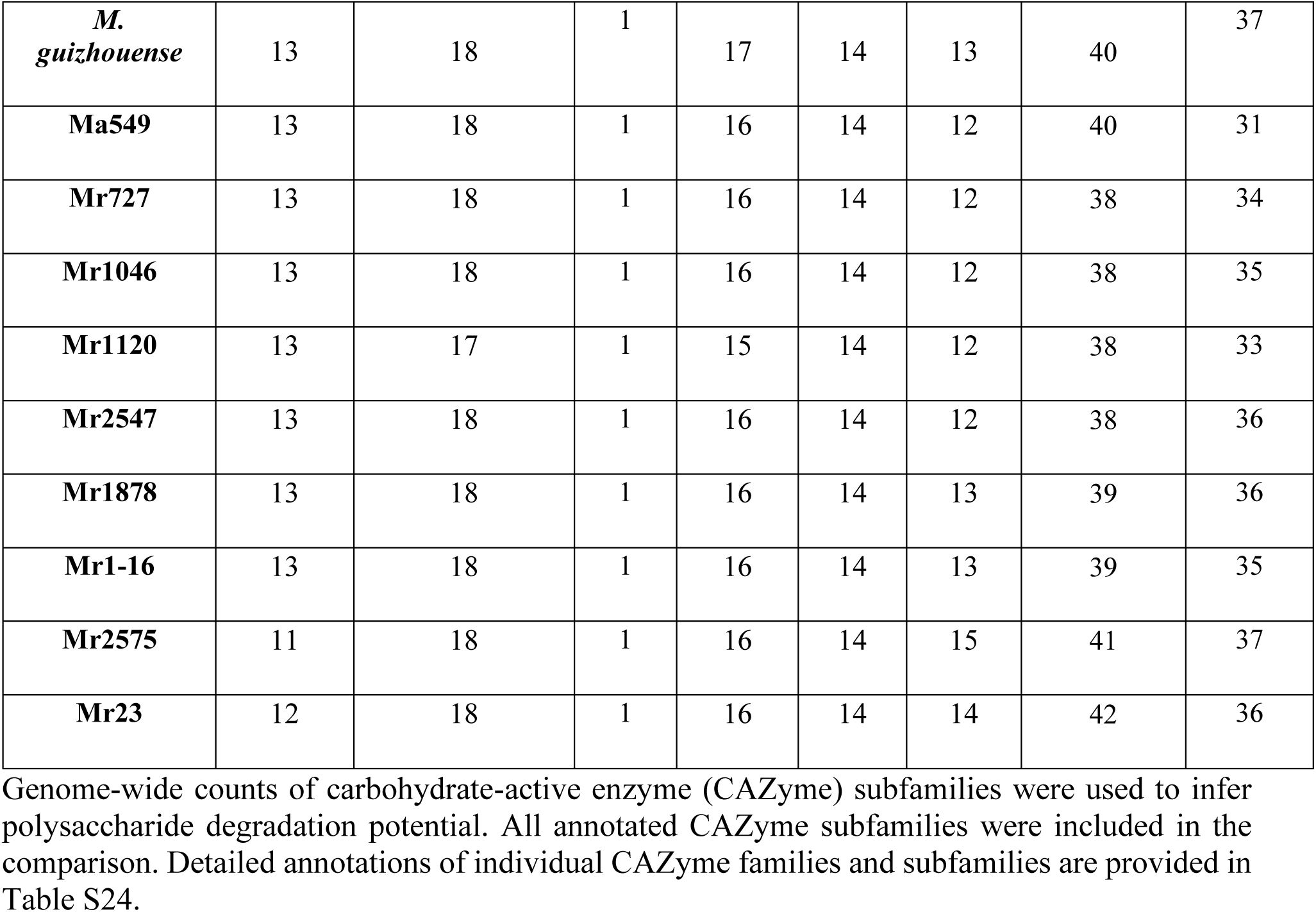
Comparative CAZyme repertoires and predicted polysaccharide degradation capacity across selected fungi.

| Species | Cellulose | Galactomannan | Inulin | Pectin | Starch | Xylan | Xyloglucan | Chitin |
| --- | --- | --- | --- | --- | --- | --- | --- | --- |
|  | GH1,<br>GH12,<br>GH5,<br>GH6,<br>GH7,<br>GH45,<br>AA9 | GH2, GH5,<br>GH26, GH27,<br>GH36, | GH32 | CE8,<br>CE12,<br>GH2,<br>GH28,<br>GH35,<br>GH43,<br>GH51,<br>GH53,<br>GH54,<br>GH78,<br>GH88,<br>GH93,<br>GH105,<br>PL1,<br>PL3,<br>PL4,<br>PL9,<br>PL11 | GH13,<br>GH15,<br>GH31 | CE1,<br>CE15,<br>GH3,<br>GH10,<br>GH11,<br>GH43,<br>GH62,<br>GH67,<br>GH115 | GH1,<br>GH3,<br>GH12,<br>GH16,<br>GH29,<br>GH31,<br>GH74,<br>GH95 | GH18,<br>GH20,<br>AA11 |
| *B. bassiana* | 11 | 17 | 1 | 15 | 13 | 17 | 46 | 29 |
| *P. chlamydosporia* | 22 | 36 | 2 | 35 | 17 | 30 | 67 | 32 |
| *M. album* | 10 | 11 | 0 | 9 | 11 | 11 | 31 | 21 |
| *M. acridum* | 14 | 17 | 2 | 10 | 13 | 11 | 36 | 23 |
| *M. majus* | 13 | 18 | 1 | 14 | 14 | 12 | 40 | 33 |
| *M. guizhouense* | 13 | 18 | 1 | 17 | 14 | 13 | 40 | 37 |
| **Ma549** | 13 | 18 | 1 | 16 | 14 | 12 | 40 | 31 |
| **Mr727** | 13 | 18 | 1 | 16 | 14 | 12 | 38 | 34 |
| **Mr1046** | 13 | 18 | 1 | 16 | 14 | 12 | 38 | 35 |
| **Mr1120** | 13 | 17 | 1 | 15 | 14 | 12 | 38 | 33 |
| **Mr2547** | 13 | 18 | 1 | 16 | 14 | 12 | 38 | 36 |
| **Mr1878** | 13 | 18 | 1 | 16 | 14 | 13 | 39 | 36 |
| **Mr1-16** | 13 | 18 | 1 | 16 | 14 | 13 | 39 | 35 |
| **Mr2575** | 11 | 18 | 1 | 16 | 14 | 15 | 41 | 37 |
| **Mr23** | 12 | 18 | 1 | 16 | 14 | 14 | 42 | 36 |

**Table 4.** Comparative auxiliary activity (AA) enzyme repertoires and predicted oxidative metabolic capacity across selected fungi.

| Species | Phe<br>noli<br>c<br>oxid<br>atio<br>n | Ligni<br>n<br>peroxi<br>dation | Alcohol/sugar<br>oxidases | Aro<br>mati<br>c<br>alco<br>hol<br>oxid<br>atio<br>n | Cop<br>per<br>radi<br>cal<br>oxi<br>das<br>es | Qui<br>non<br>e<br>redu<br>ctio<br>n | Oligosa<br>ccharid<br>e<br>oxidatio<br>n | Cytoc<br>hrom<br>e c<br>perox<br>idases | Cell<br>ulos<br>e<br>oxid<br>atio<br>n | Chit<br>in<br>oxid<br>atio<br>n | Pyra<br>nose<br>oxid<br>atio<br>n | Star<br>ch<br>oxid<br>atio<br>n | Xyl<br>an<br>oxid<br>atio<br>n |
| --- | --- | --- | --- | --- | --- | --- | --- | --- | --- | --- | --- | --- | --- |
|  | **AA**<br>**1** | **AA2** | **AA3** | **AA**<br>**4** | **AA**<br>**5** | **AA**<br>**6** | **AA7** | **AA8** | **AA**<br>**9** | **AA**<br>**11** | **AA**<br>**12** | **AA**<br>**13** | **AA**<br>**14** |
| *B. bassiana* | 11 | 3 | 13 | 2 | 3 | 1 | 23 | 2 | 1 | 4 | 2 | 0 | 0 |
| *P. chlamydosporia* | 12 | 7 | 20 | 3 | 3 | 1 | 31 | 1 | 3 | 6 | 2 | 0 | 1 |
| *M. marquandii* | 12 | 3 | 22 | 4 | 3 | 6 | 26 | 1 | 4 | 5 | 2 | 2 | 2 |
| *M. album* | 11 | 2 | 8 | 0 | 2 | 1 | 9 | 1 | 1 | 5 | 1 | 0 | 0 |
| *M. acridum* | 11 | 3 | 13 | 3 | 2 | 1 | 17 | 2 | 1 | 5 | 1 | 0 | 0 |
| *M. majus* | 15 | 3 | 18 | 2 | 2 | 1 | 27 | 2 | 1 | 6 | 2 | 0 | 1 |
| *M. guizhouense* | 14 | 3 | 22 | 3 | 2 | 1 | 31 | 0 | 1 | 6 | 2 | 0 | 1 |
| **Ma549** | 15 | 3 | 18 | 3 | 3 | 1 | 21 | 3 | 1 | 6 | 2 | 0 | 1 |
| **Mr727** | 18 | 3 | 17 | 3 | 2 | 1 | 22 | 3 | 1 | 6 | 2 | 0 | 1 |
| **Mr1046** | 18 | 3 | 18 | 2 | 2 | 1 | 26 | 3 | 1 | 6 | 2 | 0 | 1 |
| **Mr1120** | 17 | 3 | 18 | 2 | 2 | 1 | 24 | 2 | 1 | 6 | 2 | 0 | 1 |
| **Mr2547** | 18 | 3 | 17 | 2 | 2 | 1 | 26 | 3 | 1 | 6 | 2 | 0 | 1 |
| **Mr1878** | 19 | 3 | 20 | 2 | 2 | 1 | 25 | 3 | 1 | 7 | 2 | 0 | 1 |
| **Mr1-16** | 18 | 3 | 18 | 2 | 2 | 1 | 26 | 3 | 1 | 6 | 2 | 0 | 1 |
| **Mr2575** | 17 | 3 | 16 | 3 | 2 | 1 | 24 | 3 | 0 | 8 | 2 | 0 | 1 |
| **Mr23** | 18 | 3 | 18 | 3 | 2 | 1 | 25 | 3 | 0 | 7 | 2 | 0 | 1 |

Chitin-active enzymes were otherwise highly conserved, consistent with an insect-associated lifestyle. Early- and recently diverged *M. robertsii* strains encoded similar numbers of GH18 chitinases (26 vs. 27), the largest GH family in *M. robertsii*, reflecting the structural importance of chitin in arthropod exoskeletons (Elieh-Ali-Komi and Hamblin 2016). Phylogenetic analysis of GH18 genes revealed low turnover in the last 3.4 MY of *M. robertsii* evolution (Fig. S8) supporting evolutionary stability of this core functional module.

Despite the loss of processive GH6 and GH7 cellulases, all *Metarhizium* species retained a GH5_5 endo-β-1,4-glucanase and a GH12 endoglucanase. Among characterized enzymes, the *M. robertsii* GH5_5 enzyme most closely resembled Cel5b from the brown-rot fungus *Gloeophyllum trabeum* (e-value < 2.22e-126; >49% similarity) and conserves catalytic glutamate residues (Fig. S12). This retention of a non-processive cellulose-active endoglucanase despite reduced cellulolytic capacity parallels patterns described in brown-rot systems including *G. trabeum* (Floudas et al. 2012). In addition, *M. robertsii* encodes an expanded set of GH76 mannanases (13–15 genes vs. 7–9 in most fungi and ∼10 in *M. album*), suggesting increased remodeling of fungal cell walls and potential roles in morphogenetic processes such as appressorium formation.

A defining feature of the CAZyme architecture in *M. robertsii* is the enrichment of oxidative auxiliary activity (AA) enzymes. Each genome encodes ≥17 AA1 “true” laccases (phenol oxidases) (Table 4 and Table S24), exceeding numbers in other ascomycetes, including other *Metarhizium* species (Table 4), and similar in abundance to those of lignin-modifying basidiomycetes(Zerva et al. 2021). Notably, *M. robertsii* contains six to eight AA1_3 multicopper oxidases, compared to one in *P. chlamydosporia* and two in *M. marquandii*, with phylogenetic analysis suggesting *M. robertsii* -specific duplications (Fig. S13). Additional oxidoreductase families are also expanded, including AA2 lignin-modifying peroxidases, multiple AA3 subfamilies, and up to 26 AA7 glucooligosaccharide oxidases/dehydrogenases (Table S24). In contrast, saprophytic *Aspergillus* spp., known for their extensive GH repertoires, lack AA2 peroxidases and have at most 13 different laccases (in *Aspergillus niger*) (Benoit et al. 2015). This underscores a distinct metabolic strategy in *M. robertsii,* with a substantial investment in redox metabolism and reactive oxygen chemistry, that could support oxidation of phenolic and complex substrates. The AA-enriched, GH6/GH7-depleted profile closely resembles the division of labor in brown-rot fungi, where oxidative chemistry drives early substrate modification while enzymatic depolymerization capacity is reduced (Jensen et al. 2001).

Finally, AA7 enzymes catalyze the oxidation of cellooligosaccharides and chitooligosaccharides to their corresponding lactones (Haddad Momeni et al. 2021). Each *M. robertsii* strain encodes seven lactonases, including a gluconolactonase gene absent from *P. chlamydosporia*, *B. bassiana* and *M. marquandii* (Table S25). This enzyme participates in the alternative oxidative pentose phosphate pathway, and thus links CAZyme-derived oxidation products to oxidative stress responses (Jezewski et al. 2022).

### Biochemical assays confirm strong oxidative activity and limited cellulolysis

Experimental assays confirmed strong oxidative activity across *M. robertsii* strains, including oxidation of lignin-associated substrates (Fig. 6). Most strains produced brown haloes consistent with extracellular oxidative activity (Dashtban et al. 2010; Senthivelan et al. 2019) when grown on lignin or tannic acid, whereas related species (*P. chlamydosporia* and *M. marquandii*) showed little growth and activity under the same conditions (Fig. 6). *M. robertsii* strains also strongly oxidized the laccase substrate 2,2′-azino-bis(3-ethylbenzothiazoline-6-sulfonic acid) (ABTS) and, in some contexts, decolorized azure B and methylene blue (Figs. S14–S18), indicating robust redox activity whose expression was condition dependent and sometimes induced during inter-fungal confrontation (Fig. S15B). In contrast, *P. chlamydosporia* and *M. marquandii* produced clearing halos on carboxymethyl cellulose (CMC) plates confirming secreted cellulases, whereas *M. robertsii* strains did not, even with media supplemented with cellobiose that increased cellulase expression by *P. chlamydosporia* (Fig. S19). These findings support the conclusion that *M. robertsii* emphasizes oxidative chemistry rather than hydrolytic depolymerization in interactions with complex substrates.

**Fig. 6.**
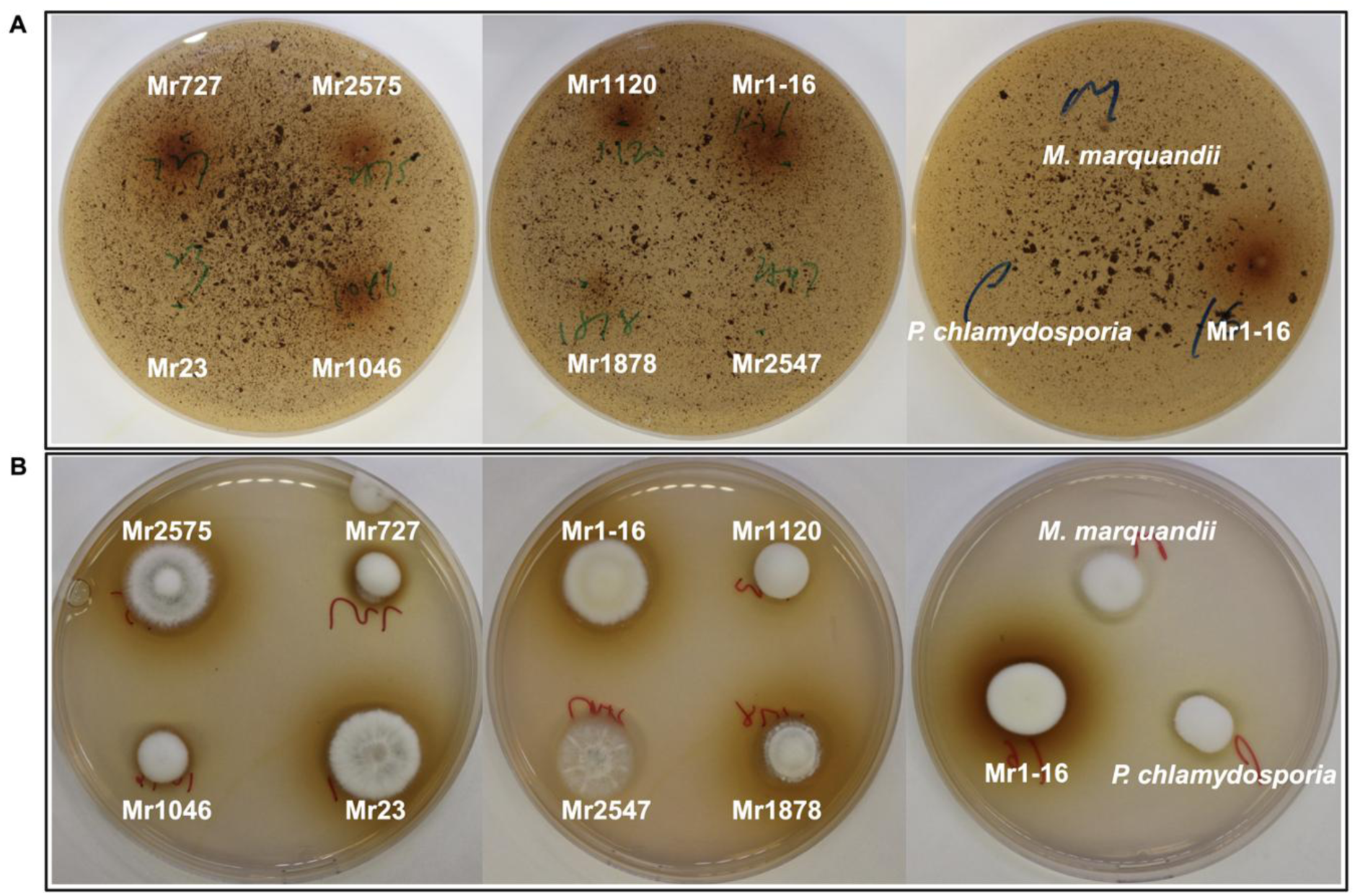
Oxidative enzyme activity of *M. robertsii* strains on lignin-associated substrates. (A) Growth on minimal medium with lignin as the sole carbon source. Most *M. robertsii* strains produced brown halos consistent with extracellular oxidative activity, whereas *P. chlamydosporia* and *M. marquandii* showed little growth or discoloration. Mr23 and Mr2547 lacked visible halo formation. (B) Growth on potato dextrose agar supplemented with 0.1% tannic acid. Brown halo formation indicative of laccase-associated oxidation was observed in all *M. robertsii* strains except Mr2547, while comparator species showed no discoloration. Reverse plate images are shown in Fig. S14.

### Regulatory variation and secondary metabolism contribute to phenotypic divergence

Secondary metabolites (SMs) contribute to ecological specialization (Fox and Howlett 2008), and *Metarhizium* spp. are prolific producers (St. Leger 2024). The total number of predicted biosynthetic gene clusters (BGCs) varied between strains, with early diverged lineages encoding significantly more BGCs than recently diverged strains (102 vs. 92, p = 0.006) (Table 5). This difference was primarily driven by type I polyketide synthase (T1PKS) and nonribosomal peptide synthetase (NRPS) classes (26 vs. 22 T1PKS; 19 vs. 15 NRPS), whereas terpene BGC counts remained constant between early and recently diverged strains (11 vs. 11) indicating selective diversification of specific SM pathways rather than global expansion.

**Table 5.**
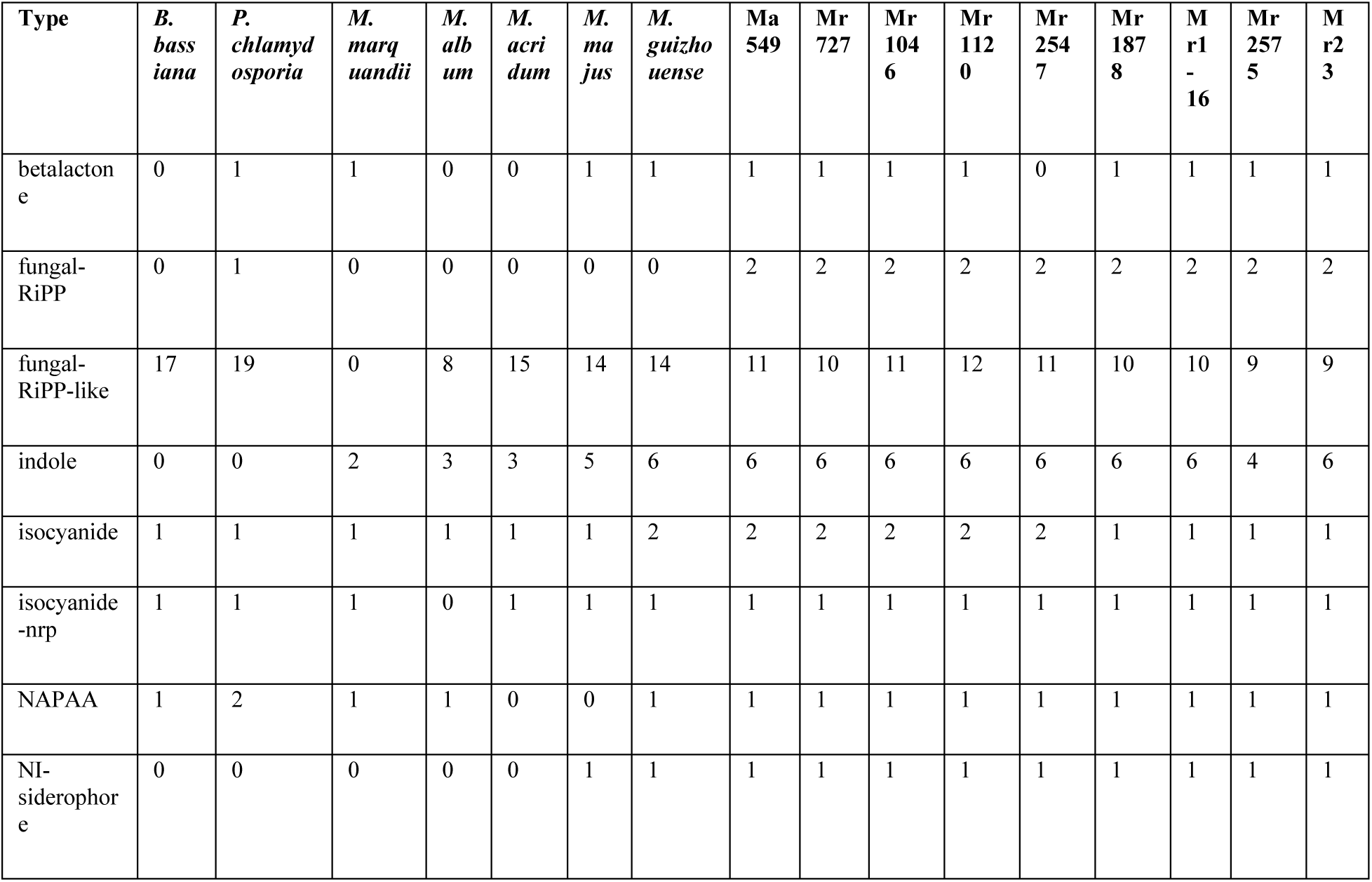
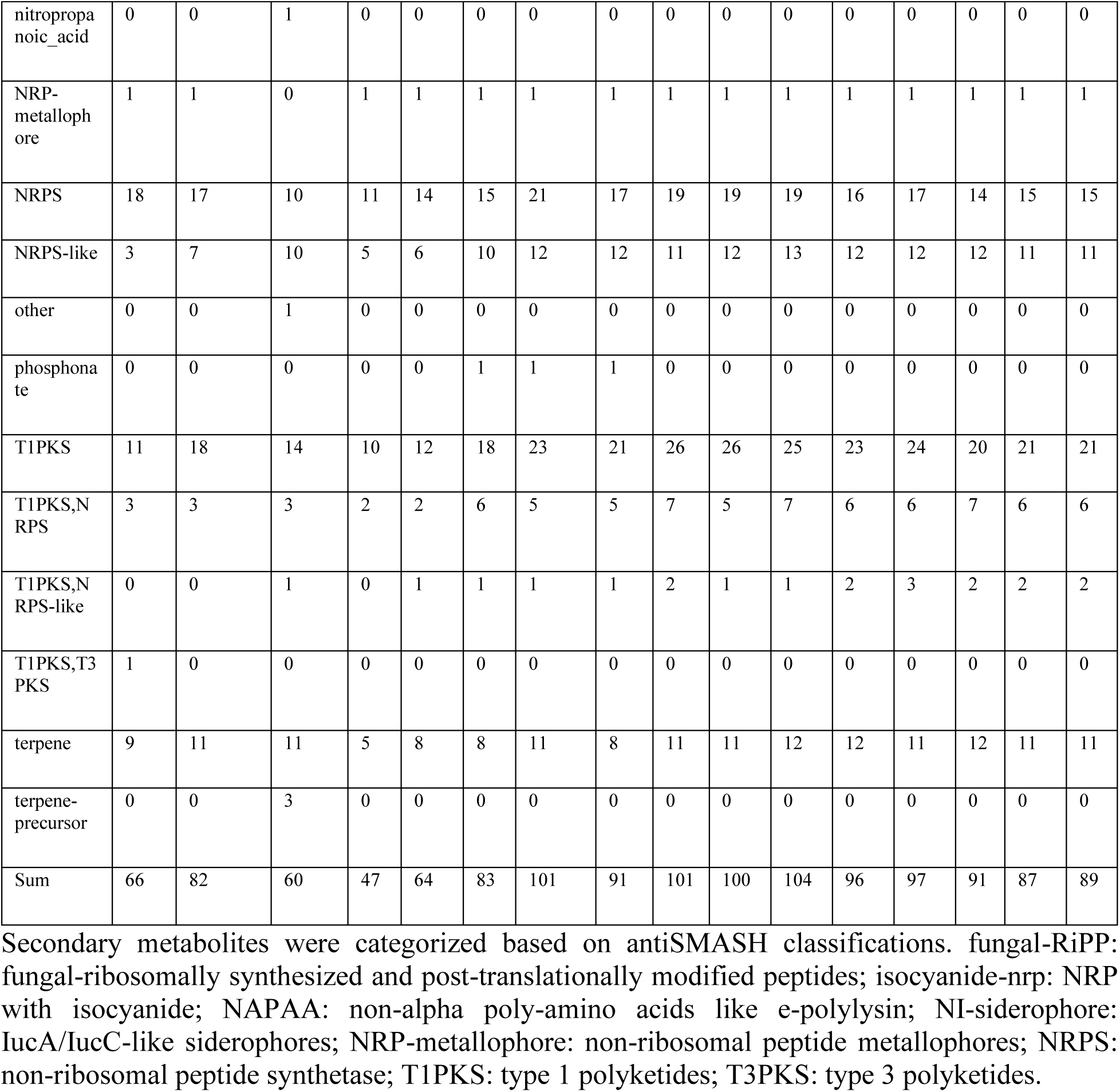
Distribution of predicted secondary metabolite biosynthetic gene clusters across fungal genomes.

| Type | *B. bassiana* | *P. chlamydosporia* | *M. marquandii* | *M. albidum* | *M. acridum* | *M. majus* | *M. guizhouense* | Ma 549 | Mr 727 | Mr 1046 | Mr 1120 | Mr 2547 | Mr 1878 | Mr 1 - 16 | Mr 2575 | Mr 23 |
| --- | --- | --- | --- | --- | --- | --- | --- | --- | --- | --- | --- | --- | --- | --- | --- | --- |
| betalactone | 0 | 1 | 1 | 0 | 0 | 1 | 1 | 1 | 1 | 1 | 1 | 0 | 1 | 1 | 1 | 1 |
| fungal-RiPP | 0 | 1 | 0 | 0 | 0 | 0 | 0 | 2 | 2 | 2 | 2 | 2 | 2 | 2 | 2 | 2 |
| fungal-RiPP-like | 17 | 19 | 0 | 8 | 15 | 14 | 14 | 11 | 10 | 11 | 12 | 11 | 10 | 10 | 9 | 9 |
| indole | 0 | 0 | 2 | 3 | 3 | 5 | 6 | 6 | 6 | 6 | 6 | 6 | 6 | 6 | 4 | 6 |
| isocyanide | 1 | 1 | 1 | 1 | 1 | 1 | 2 | 2 | 2 | 2 | 2 | 2 | 1 | 1 | 1 | 1 |
| isocyanide-nrp | 1 | 1 | 1 | 0 | 1 | 1 | 1 | 1 | 1 | 1 | 1 | 1 | 1 | 1 | 1 | 1 |
| NAPAA | 1 | 2 | 1 | 1 | 0 | 0 | 1 | 1 | 1 | 1 | 1 | 1 | 1 | 1 | 1 | 1 |
| NI-siderophore | 0 | 0 | 0 | 0 | 0 | 1 | 1 | 1 | 1 | 1 | 1 | 1 | 1 | 1 | 1 | 1 |
| nitropropionic_acid | 0 | 0 | 1 | 0 | 0 | 0 | 0 | 0 | 0 | 0 | 0 | 0 | 0 | 0 | 0 | 0 |
| NRP-metallophore | 1 | 1 | 0 | 1 | 1 | 1 | 1 | 1 | 1 | 1 | 1 | 1 | 1 | 1 | 1 | 1 |
| NRPS | 18 | 17 | 10 | 11 | 14 | 15 | 21 | 17 | 19 | 19 | 19 | 16 | 17 | 14 | 15 | 15 |
| NRPS-like | 3 | 7 | 10 | 5 | 6 | 10 | 12 | 12 | 11 | 12 | 13 | 12 | 12 | 12 | 11 | 11 |
| other | 0 | 0 | 1 | 0 | 0 | 0 | 0 | 0 | 0 | 0 | 0 | 0 | 0 | 0 | 0 | 0 |
| phosphonate | 0 | 0 | 0 | 0 | 0 | 1 | 1 | 1 | 0 | 0 | 0 | 0 | 0 | 0 | 0 | 0 |
| T1PKS | 11 | 18 | 14 | 10 | 12 | 18 | 23 | 21 | 26 | 26 | 25 | 23 | 24 | 20 | 21 | 21 |
| T1PKS,NRPS | 3 | 3 | 3 | 2 | 2 | 6 | 5 | 5 | 7 | 5 | 7 | 6 | 6 | 7 | 6 | 6 |
| T1PKS,NRPS-like | 0 | 0 | 1 | 0 | 1 | 1 | 1 | 1 | 2 | 1 | 1 | 2 | 3 | 2 | 2 | 2 |
| T1PKS,T3PKS | 1 | 0 | 0 | 0 | 0 | 0 | 0 | 0 | 0 | 0 | 0 | 0 | 0 | 0 | 0 | 0 |
| terpene | 9 | 11 | 11 | 5 | 8 | 8 | 11 | 8 | 11 | 11 | 12 | 12 | 11 | 12 | 11 | 11 |
| terpene-precursor | 0 | 0 | 3 | 0 | 0 | 0 | 0 | 0 | 0 | 0 | 0 | 0 | 0 | 0 | 0 | 0 |
| Sum | 66 | 82 | 60 | 47 | 64 | 83 | 101 | 91 | 101 | 100 | 104 | 96 | 97 | 91 | 87 | 89 |

Secondary metabolites were categorized based on antiSMASH classifications. fungal-RiPP: fungal-ribosomally synthesized and post-translationally modified peptides; isocyanide-nrp: NRP with isocyanide; NAPAA: non-alpha poly-amino acids like e-polylysine; NI-siderophore: IucA/IucC-like siderophores; NRP-metallophore: non-ribosomal peptide metallophores; NRPS: non-ribosomal peptide synthetase; T1PKS: type 1 polyketides; T3PKS: type 3 polyketides.

Despite quantitative differences, many BGCs were conserved across strains, defining a substantial shared “core” SM repertoire. All strains encoded clusters associated with destruxins (dtx), serinocyclins, metachelin, ochratoxin A, cytochalasin E/cytochalasin K, BAB/BAA polyketides, subglutinol A, terpendole E, NG-391 (a highly mutagenic 7-desmethyl analog of fusarin C(Bohnert et al. 2013)), and various antimicrobial compounds (e.g., ustilaginoidin, pseurotin, viridicatum, helvolic acid) (Table S26), consistent with roles in competitive interactions across soil, insect, and rhizosphere environments.

Importantly, variation in cluster organization, partial retention, and likely expression differences indicate that phenotypic diversity arises not from wholesale gain or loss of pathways, but from structural and regulatory modulation of a conserved biosynthetic framework. At the genomic level, BGC organization is dynamic. For example, the subglutinol BGC illustrates partial retention and lineage-specific decay of an ancestrally acquired pathway (Table S27). The complete six-gene cluster (subA–F), previously inferred to have entered *Metarhizium* via horizontal transfer (Lin et al. 2014; Kato et al. 2016), was retained in only a subset of strains (Mr23, Mr2575, *M. acridum*), whereas others retained incomplete gene sets with variable sequence conservation (Table S27). Given that partial synteny often yields structurally related but distinct metabolites, these truncated clusters likely encode divergent products rather than inactive remnants.

Other examples of loss include the pseurotin BGC, previously identified in *M. anisopliae* E6 that was absent from *M. anisopliae* Ma549. A beta-lactone BGC, also associated with antimicrobial activity(Robinson et al. 2019), was present in all strains except Mr2547, indicating a recent strain-specific loss (Table 5). Iron acquisition represents a conserved and potentially critical ecological function. All *M. robertsii* strains along with Ma549, *M. majus*, and *M. guizhouense,* encoded an aerobactin-like siderophore BGC that was absent from *M. album*, *M. acridum*, and *P. chlamydosporia* (Table 5). The core gene of this cluster most closely resembled a siderophore gene from the soil endophyte *Dichotomopilus funicola* (E-value: 0; 57% identity) suggesting a conserved role in iron acquisition within plant- and insect-associated niches. In addition, all *M. robertsii* strains possessed two siderophore-associated clusters linked to mrsidA and mrsidD involved in iron homeostasis in Mr23(Zhang et al. 2021). While mrsidA was identical across strains (Table S28), mrsidD exhibited sequence divergence, particularly in early diverged lineages (Table S29), indicating ongoing fine-scale evolution of iron acquisition pathways.

Although some strains (e.g., Mr727, Mr1046, Ma549) do not produce destruxins at toxic levels in culture (Wang et al. 2023; Sheng and St. Leger 2026), the previously identified core Mr23 destruxin locus (dtxS1–dtxS4) (Wang et al. 2012) is highly conserved across strains (>98% identity for *dtxS1* and *dtxS2*) (Fig. S20; Table S30). Structural variation is nonetheless evident: Mr23 and Mr2575 retain separate *dtxS3* and *dtxS4* genes, whereas other strains carry a fused *dtxS3–dtxS4* gene (Fig. S20). Mr2575 also lacked several genes upstream of *dtxS1* that are present in other strains, but disruption of upstream genes does not affect destruxin production in Mr23 (Wang et al. 2012). Collectively, these observations support the conclusion that strain differences in destruxin output are primarily driven by regulatory changes and local locus restructuring rather than loss of core biosynthetic genes. Together, these results mirror patterns observed for CAZymes and proteases: a conserved core functional repertoire is maintained across *M. robertsii*, while strain-level diversification arises through targeted gene cluster gain and loss, structural rearrangement, and regulatory divergence.

### Signal transduction and regulatory gene repertoires

Signal transduction and regulatory gene repertoires were also conserved in overall composition across *M. robertsii* strains. Compared to *M. anisopliae* 549, *M. robertsii* strains carried approximately ten additional G protein-coupled receptors (GPCRs), primarily within the PTH11-like family, suggesting enhanced capacity for sensing environmental cues. However, GPCR counts were similar between early and recently diverged *M. robertsii* strains (Tables S31–S32), and most receptor families were represented by single orthologs, indicating that this expansion predates intraspecific diversification.

Mr2575 exhibited two recent GPCR duplications, including one PTH11-like receptor and one membrane progesterone receptor (mPR)-like receptor, and also contained a positively selected PTH11-like GPCR, suggesting lineage-specific tuning of environmental sensing. In addition, a single GprK homolog in Mr1120 was under positive selection. Beyond these cases, most GPCRs showed neither duplication nor signatures of selection, highlighting strong conservation of receptor repertoires across strains.

Transcription factor (TF) inventories were similarly stable with *M. robertsii* strains encoding an average of 448 TFs and showing minimal differences between early and recently diverged strains (442 vs. 451; Table S33). Despite overall conservation, evidence of adaptive divergence was found within regulatory networks. Notably, 25 Zn2Cys6 transcription factors (ZnFTFs) were present in only a subset of strains, with a phylogeny suggesting that this was due to duplication events followed by lineage specific gene loss rather than recent horizontal acquisition (Fig S21). Two of these duplicated ZnFTFs showed weak similarity to characterized regulators, including Cutinase Transcription Factor 1 from *Fusarium solani* (20.1% identity) and the bikaverin cluster regulator Bik5 from *F. fujikuroi* (16.1%), limiting functional inference. Among 11 ZnFTFs under positive selection, sequence similarity to experimentally characterized regulators was likewise modest, ranging from ∼12–49% identity (Tables S34-35).

Core developmental regulators were highly conserved. The central asexual sporulation pathway genes were present as single copy orthologs in all strains (Table S36). Mating-type distribution followed a heterothallic pattern, with strains carrying either MAT1-1-1 or MAT1-2-1 idiomorphs. Homologs of pheromone signaling and ascospore-associated genes were also conserved across strains (Table S36), suggesting retention of the genetic capacity for sexual reproduction despite the absence of observed teleomorphs.

### Dynamic Non-coding RNA Systems

Non-coding RNA repertoires showed substantially greater variability across *M. robertsii* strains than most protein-coding functional classes. Transfer RNA (tRNA) gene complements were reduced relative to *M. anisopliae* Ma549 (160 genes), with all *M. robertsii* genomes encoding fewer copies (mean 127; Table S37). Mr23 retained the highest number among the *M. robertsii* strains (142), whereas other lineages exhibited further reductions, indicating lineage-specific contraction of tRNA copy number. Suppressor tRNAs were unevenly distributed: all strains except Mr727 and Mr1120 encoded a predicted nonsense suppressor tRNA recognizing the UAG stop codon, while Mr23 uniquely carried a predicted pseudo-suppressor tRNA recognizing UGA. These patterns suggest differential retention or loss of translational readthrough.

Predicted microRNA (miRNA) repertoires were even more dynamic. Total miRNA counts ranged from 26 in Mr1-16 to 55 in Mr23 and spanned 67 families (Tables S37–S38), with only seven families conserved across all strains. Most families were lineage-restricted, indicating rapid turnover of small RNA complements within *M. robertsii*. For example, Mr727 encoded members of 35 miRNA families, whereas Mr1-16 contained only 18, highlighting substantial heterogeneity even among closely related strains. One family (MIR827_2) was detected exclusively in recently diverged strains. MIR827 homologs are associated with phosphate regulation in plants (Hackenberg et al. 2013), but the *M. robertsii* sequences shared only moderate similarity (47–57%) with these leaving their evolutionary origin and functional significance unresolved.

These results indicate that non-coding RNA repertoires, particularly miRNAs, are among the most dynamic components of the *M. robertsii* genome. This contrasts with the stability of core metabolic and regulatory gene inventories, and parallels the strain-specific patterns observed for transposable elements and RIP signatures. Collectively, these features point to ongoing diversification of genome defense and post-transcriptional regulatory networks as a key axis of intraspecific variation.

## Discussion

Our comparative analysis demonstrates that substantial ecological divergence—from slower-killing, highly sporulating strains to faster-killing lineages with enhanced endophytic capacity (Sheng and St. Leger 2026), can arise over short evolutionary timescales without extensive genome-wide innovation. Rather than requiring wholesale genetic restructuring, diversification proceeds through targeted genomic and regulatory modifications within a highly conserved framework. These findings support a hierarchical model of genome evolution in which a conserved core genome encodes the fundamental entomopathogenic lifestyle, while diversification emerges from dynamic peripheral layers. These peripheral systems include repeat-rich genomic compartments shaped by transposable elements and RIP, regulatory networks subject to duplication and selection, and post-transcriptional systems including rapidly evolving non-coding RNAs. Together, these layers enable fine-scale functional tuning and ecological adaptation without requiring large-scale innovation in gene content or genome structure.

A defining feature of *M. robertsii* evolution is the exceptional conservation of genome architecture. High nucleotide identity, extensive macrosynteny, and preservation of gene order—even across deep evolutionary timescales within *Metarhizium*—indicate strong constraints on large-scale genomic reorganization. Core functional repertoires, including virulence-associated genes, proteases, carbohydrate-active enzymes, and central developmental regulators, remain similarly stable across strains. These observations support the view that the fundamental entomopathogenic toolkit is encoded within a conserved genomic backbone. This pattern aligns with previous comparative analyses that identified shared “entomopathogenicity toolkits” conserved across deeply divergent lineages(Gao et al. 2011; Hu et al. 2014; St. Leger and Wang 2020).

In stark contrast to this structural stability, genomic divergence involves repeat-rich regions characterized by variable transposable element (TE) content, repeat-induced point mutation (RIP) activity, and localized synteny disruption. Critically, these dynamic regions are largely shared across strains rather than confined to lineage-specific chromosomes, indicating that diversification occurs within a common genomic substrate rather than through wholesale acquisition of novel compartments. Although accessory chromosomes have been described in *Metarhizium*(Habig et al. 2024), their fragmented and redistributed representation across *M. robertsii* genomes suggests that they contribute to, but do not solely define, genomic plasticity. This fragmentation and integration pattern may represent a mechanism by which fungi can “trial” new genetic material before potentially stabilizing beneficial elements within the core genome (Croll and McDonald 2012).

Within these dynamic compartments, multiple evolutionary processes act synergistically to generate localized genomic innovation. TE proliferation strongly associates with increased gene duplication, supporting a central role for mobile DNA in promoting structural variation and gene family diversification (Castanera et al. 2016; Faino et al. 2016). Strains display markedly different RIP signatures, likely reflecting variation in genome defense efficacy, recent sexual history, or lineage-specific regulation of mutational processes. RIP activity further modifies duplicated sequences, introducing mutations that accelerate functional divergence while simultaneously constraining TE expansion (Gladyshev 2017; Van Wyk et al. 2021). Together, these mechanisms establish a dynamic equilibrium between innovation and genome stability: TEs generate variation, RIP shapes and limits this variation, and both processes remain spatially restricted within the genome. These dynamic regions correspond to known evolutionary hotspots, particularly subtelomeric compartments that are characterized by elevated rates of recombination, repeat accumulation, and chromosomal rearrangements across diverse fungal lineages. However, the spatial distribution of repeat content varies substantially among strains, with some (Mr2575, Mr23) showing relatively low subtelomeric sequence content (∼5%) despite high overall TE loads, suggesting that repeat expansion can occur through alternative mechanisms that bypass traditional subtelomeric accumulation patterns. Thus, TE dynamics, RIP activity, and fragmented accessory chromosome-derived sequences collectively define a shared set of genomic regions that undergo repeated modification across strains without destabilizing core genome function.

Overlaying this structural framework, regulatory and post-transcriptional evolution emerge as primary drivers of phenotypic divergence. Although transcription factor repertoires and signaling components remain largely conserved in copy number, lineage-specific duplications and signatures of positive selection indicate fine-scale tuning of regulatory networks. The enrichment of positively selected genes in transporter systems, signaling pathways, and intracellular regulatory processes suggests that adaptation proceeds primarily through modification of gene function and expression rather than through acquisition of entirely novel gene families. This pattern aligns with broader evolutionary theory emphasizing regulatory change as a major driver of phenotypic diversification (Carroll 2008). Strain Mr2575 exemplifies this pattern, showing particular enrichment of positively selected genes and recent duplications while maintaining similar total gene counts to related strains. For instance, Mr2575 and Mr23 encode similar numbers of trypsins, yet Mr2575 exhibits more gain and loss events than Mr23 relative to Mr1-16 (Fig. S10), consistent with an accelerated tempo of gene turnover rather than net expansion.

Non-coding RNA systems represent a particularly dynamic layer of post-transcriptional evolution. In contrast to the remarkable stability of protein-coding gene inventories, extensive lineage-specific gain and loss of miRNA families, combined with variation in suppressor tRNAs, indicates rapid turnover of post-transcriptional regulatory systems. This variability parallels patterns observed in transposable element content and RIP activity, suggesting functional coupling between genome defense processes and RNA-mediated regulation.

Metabolic evolution in *M. robertsii* illustrates the primacy of functional repurposing over gene acquisition in ecological adaptation. Despite significant ecological divergence—with recently diverged strains becoming broad host range plant associates capable of utilizing diverse plant sugars (Sheng and St. Leger 2026)—core enzymatic repertoires, including proteases and carbohydrate-active enzymes, remain highly conserved across strains. Instead of expanding canonical plant cell wall-degrading systems, *M. robertsii* exhibits a conserved reduction in processive cellulases alongside expansion of oxidative auxiliary activity enzymes. This enzymatic configuration produces a metabolic profile convergent with brown-rot fungi, in which oxidative chemistry drives substrate modification while enzymatic depolymerization capacity is reduced (Floudas et al. 2012). However, in *M. robertsii*, this oxidative capacity appears repurposed for alternative ecological functions, including detoxification of insect-derived phenolic compounds and specialized rhizosphere interactions. For example, Mr2575 produces hydrogen peroxide in the presence of quinones to mount non-enzymatic attacks on insect cuticular melanin(St. Leger et al. 1988). This convergence with brown-rot fungi therefore highlights how similar biochemical strategies can evolve independently to serve distinct ecological roles.

Secondary metabolite biosynthesis provides additional evidence for the importance of regulatory evolution over structural innovation. Although most biosynthetic gene clusters remain conserved across strains, variation in cluster organization, partial gene retention, and regulatory output generates substantial phenotypic diversity. The destruxin biosynthetic cluster exemplifies this pattern: non-toxigenic early diverged *Metarhizium* species such as *M. acridum* lack this cluster entirely (Amiri-Besheli et al. 2000; Wang et al. 2004), which likely entered the PARB clade (*M. pinghaense*, *M. anisopliae*, *M. robertsii*, and *M. brunneum*) through horizontal transfer and contributed to broad host range evolution(Zhang et al. 2019). Despite retaining intact destruxin clusters, strains Mr1046 and Mr727 exhibit non-toxigenic phenotypes and proliferate within hosts similarly to *M. acridum* (Sheng and St. Leger 2026). Thus, differences in destruxin synthesis likely reflect regulatory divergence and local structural variation rather than presence-absence polymorphisms. This pattern reinforces the broader conclusion that ecological differentiation proceeds through modulation of existing functional capacity rather than wholesale expansion of metabolic repertoires.

An unexpected finding was the heterogeneous distribution of subtelomeric sequence content across strains, with some highly TE-active strains (Mr2575, Mr23) showing relatively low subtelomeric content (∼5%) compared to other strains (∼20%). This pattern suggests that the mechanisms driving repeat expansion are not uniformly distributed across chromosomal regions and that high overall TE activity does not necessarily translate to subtelomeric accumulation. While this discrepancy could potentially reflect technical artifacts related to assembly methodology differences between PacBio-Illumina hybrid and Illumina-only approaches, it may also indicate genuine biological variation in the spatial dynamics of TE insertion and proliferation. Such strain-specific differences in repeat distribution patterns could contribute to the localized genomic innovation that enables rapid ecological adaptation while maintaining overall genome stability.

These results collectively support a hierarchical, multi-layer model of genome evolution in which ecological diversification emerges from the coordinated interaction of distinct but interconnected genomic processes. At the foundation, a highly conserved core genome encodes the fundamental biological capacity of the organism. Surrounding this stable core, repeat-rich genomic compartments generate structural variation through TE activity and gene duplication, while genome defense mechanisms such as RIP shape and constrain this variation. Regulatory networks, including transcription factors and signaling pathways, provide fine-scale control over gene expression, while rapidly evolving non-coding RNA systems add an additional layer of post-transcriptional regulation. The integration of these processes enables rapid ecological adaptation without requiring large-scale changes in gene content or genome architecture, reconciling strong genomic constraint with substantial phenotypic flexibility.

This layered evolutionary model may represent a general mechanism underlying rapid ecological divergence in eukaryotic microbes, particularly those facing complex and variable environmental challenges. By channeling innovation through specific genomic processes while preserving core functional architecture, organisms can achieve adaptive flexibility while maintaining the genetic coherence necessary for complex life cycles. Understanding these hierarchical evolutionary processes has important implications for predicting microbial responses to environmental change and for harnessing genomic diversity in biotechnological applications. The rapid evolutionary rates documented here (TE proliferation at ∼43 elements/MY, RIP activity at 0.9–13.8%/MY) suggest that environmental challenges may trigger accelerated genome evolution in timeframes relevant to climate change and anthropogenic disturbances. Furthermore, the modular architecture identified in this system provides a framework for engineering enhanced microbial biocontrol agents by leveraging natural variation in repeat dynamics, regulatory networks, and metabolic repurposing rather than relying solely on transgenic approaches.

## Methods

### Fungal strains and culture conditions

Seven *M. robertsii* strains (ARSEF Mr1046, Mr1120, Mr1878, Mr23, Mr2547, Mr2575, and Mr727) were obtained from the USDA Entomopathogenic Fungus Collection (Ithaca, NY, USA). Mr1-16 was isolated from rhizospheric soil collected in Burkina Faso by Dr. Etienne Bilgo. Strains were revived from −80°C stocks 12 days before each experiment and maintained on potato dextrose agar (PDA) at 27°C.

### Plate assays for cellulolytic and oxidative enzyme activity

Extracellular cellulase activity was evaluated on minimal salts agar (1.5% Noble agar, 0.2% NaNO_₃_, 0.1% KH_₂_PO_₄_, 0.05% MgSO_₄_, FeSO_₄_, ZnSO_₄_, CuSO_₄_, and MnCl_₂_ each at 10 mg per 100 mL) supplemented with 1% (w/v) carboxymethylcellulose sodium salt (CMC) as the sole carbon source. Laccase-mediated oxidation was assessed using ABTS (2,2′-azino-bis[3-ethylbenzothiazoline-6-sulfonic acid]) and tannic acid on PDA supplemented with 0.1% (w/v) substrate, and using lignin (alkali lignin; Sigma, CAS 8068-05-1, product reference 471003) either as 1% (w/v) sole carbon source in minimal salts medium or as an additive to PDA. Oxidation of tannic acid or lignin produces brown pigmentation (Sharma et al. 2017; Buzzo et al. 2024), and ABTS oxidation produces a blue-green product that can darken to purple over time (Fernández-Remacha et al. 2022). To screen for lignin peroxidase activity, Azure B (0.01% w/v) was included in agar media; decolorization in the presence of H_₂_O_₂_ is indicative of Lignin peroxidase activity (Archibald 1992). Methylene blue (0.01% w/v) was also incorporated into PDA to assess redox-associated dye decolorization. ABTS, Azure B, tannic acid and methylene blue were sterilized by filtration and added to autoclaved media after cooling.

### Genome sequencing, assembly, and gene annotation

Single-spore isolates were generated for Mr727, Mr1046, Mr1120, Mr1878, Mr2547, and Mr1-16 and used for DNA extraction and Illumina HiSeq2500 sequencing. Mr2575 was sequenced using PacBio RSII and Illumina NovaSeq 6000 (NOVOGENE). Genome assemblies were generated using SOAPdenovo (Luo et al. 2012) with exploration of multiple k-mers, followed by gap filling and assembly optimization using krskgf and GapCloser. Gene models were predicted using AUGUSTUS (Stanke et al. 2004). Assembly completeness was assessed with BUSCO using the fungal lineage dataset (Simão et al. 2015), and N50 statistics were computed using the R package CNEr (Tan et al. 2019). Assembly quality metrics revealed substantial differences between sequencing approaches: the PacBio-Illumina hybrid assembly of Mr2575 achieved superior contiguity (N50: 3.2 Mb, 12 contigs) compared to Illumina-only assemblies (N50: 0.15–0.8 Mb, 45–180 scaffolds), but showed different patterns of repeat region resolution. Long-read sequencing enabled span of most repetitive regions in Mr2575, potentially leading to collapse of highly similar subtelomeric repeats into single consensus sequences, whereas Illumina assemblies fragment at repetitive boundaries, preserving individual repeat copies as separate contigs. This methodological difference could partially explain the observed subtelomeric content discrepancy (∼5% in Mr2575 vs. ∼20% in Illumina-assembled genomes). tRNA and rRNA genes were predicted using tRNAscan-SE (Chan and Lowe 2019) and rRNAmmer (Lagesen et al. 2007), respectively. Other non-coding RNAs (including miRNAs, sRNAs, and snRNAs) were predicted by searches against Rfam using cmsearch (Cui et al. 2016; Kalvari et al. 2018).

### Whole-genome comparison and synteny

Average nucleotide identity (ANI) was calculated for pairwise comparisons between *M. robertsii* and *M. anisopliae* ARSEF549 (Ma549) using an ANI calculator (Rodriguez-R and Konstantinidis 2014). Whole-genome alignments among *M. robertsii* strains and Ma549 were performed using MUMmer v4.0.0 (Marçais et al. 2018) with NUCmer and default parameters. Dot plots for visualization and downstream analyses were generated in R using GenomicRanges (Lawrence et al. 2013; Wickham et al. 2019; Iwanicki et al. 2022).

### Accessory chromosome–derived sequence identification

Accessory chromosome–derived sequences were identified by aligning genome assemblies to chrA and chrB from the R3-I4 strain using NUCmer (MUMmer v4) with a minimum identity of 85% and alignment length ≥1 kb(Marçais et al. 2018; Habig et al. 2024). Alignment coordinates were extracted using show-coords, and reference intervals were merged to obtain non-overlapping regions. Alignment coverage was calculated as the proportion of the reference chromosome covered by unique homologous alignments. The number of alignment blocks and scaffolds containing homologous regions was used to assess fragmentation.

### Subtelomeric enrichment analysis

Synteny gap intervals from pairwise genome comparisons were analyzed in the coordinate system of the anchor genome. Subtelomeric regions were defined as 100 kb windows flanking predicted telomeric repeat tracts (TTAGGG/CCCTAA), with scaffold ends considered telomere-associated if ≥5 repeat copies were detected within the terminal 20 kb. Gap intervals were intersected with subtelomeric regions using BEDTools. Enrichment was assessed by comparing the proportion of gap bases located in subtelomeric regions to the genome-wide subtelomeric fraction using Fisher’s exact test.

To evaluate potential assembly bias, we additionally quantified the proportion of each genome located within 100 kb of scaffold ends independent of telomere detection.

### Repetitive element annotation and RIP analyses

Repetitive sequences were identified in *Metarhizium* spp. and comparator fungi (*Pochonia chlamydosporia*, *Cordyceps militaris*, *Neurospora crassa*, *Beauveria bassiana*, and *Epichloë festucae*) using a combined de novo and homology-based strategy. Genomes were aligned using PALS, repetitive regions were identified with PILER (Edgar and Myers 2005), and results were combined with RepeatScout v1.0.5 to generate repeat families (Price et al. 2005). RepeatMasker v4.1.6 was then used for repeat annotation with Repbase v29.05 restricted to the “fungi” taxon (Bao et al. 2015). Subtelomeric sequence content was estimated by quantifying repetitive DNA sequences within 50 kb of chromosome termini, with terminal regions identified through analysis of telomeric repeat motifs and assembly boundaries.

To assess conservation of RIP-associated machinery in *M. robertsii*, BLASTp searches (E-value ≤ 1 e-5) were performed using *N. crassa* RIP pathway proteins, including RID and DIM-2 and cofactors implicated in DIM-2–mediated DNA methylation and heterochromatin formation (HP-1, DDB-1, Cul4, DIM-5, and DIM-7) (Lewis et al. 2010; Gessaman and Selker 2017; He et al. 2020). Similarity among homologous protein sets was evaluated by multiple sequence alignment using Clustal Omega v1.2.4 (Sievers et al. 2011).

Genome-wide RIP signatures were quantified using RIPCAL v1.0 and The RIPper (Hane and Oliver 2008; Van Wyk et al. 2019). Analyses used 1,000 bp sliding windows with 500 bp steps. The RIPper was additionally used to identify large RIP-affected regions (LRARs) and to compute RIP composite indices and GC content using stringent thresholds (RIP product > 1.15, RIP substrate ≤ 0.75, and composite index > 0). *N. crassa* served as a positive control.

### Orthology inference and phylogenomic analysis

Orthologs and gene duplication events among the eight *M. robertsii* strains and Ma549 were inferred with OrthoFinder v2.5.2(Emms and Kelly 2019). Genes were classified as core (present in all strains), shared (present in a subset), or strain-specific (unique to one strain), following standard comparative-genomic conventions. Protein sequences used for gene-tree reconstruction were aligned with Clustal Omega v1.2.4(Sievers et al. 2011). The best-fitting substitution model was selected using a RAxML-NG v1.2.1 Perl script(Kozlov et al. 2019), and maximum-likelihood trees were reconstructed with 100 bootstrap replicates.

### Functional annotation and gene family analyses

Protein domains and families were annotated using InterProScan v5.50-84.0 with Pfam 33.1(Jones et al. 2014; Mistry et al. 2021). For broader functional annotation, proteomes were searched against multiple databases using BLASTp(Camacho et al. 2009). CAZymes were identified via dbCAN3(Cantarel et al. 2009; Zheng et al. 2023), and biosynthetic gene clusters were predicted using fungal antiSMASH v7.1.0(Medema et al. 2011). Cytochrome P450s were extracted using Pfam PF00067 and classified against CYPED v6.0 (E-value ≤ 1e−5) (Sirim et al. 2009). Signal peptides and transmembrane domains were predicted using SignalP v6.0 and Phobius, respectively (Käll et al. 2004; Kall et al. 2007; Almagro Armenteros et al. 2019). Transporters and channels were annotated using TCDB(Saier et al. 2021). Virulence-associated genes were identified by homology to PHI-base using BLASTp (E-value ≤ 1e−20) (Urban et al. 2019); PHI entries annotated as not affecting pathogenicity were excluded from downstream analyses. Kinases were classified by BLASTp against KinBase (http://kinase.com). Candidate GPCRs were identified using family-specific HMMs constructed from *Magnaporthe grisea* GPCRs(Li et al. 2007); HMMER searches used E-value < 1e−5(Eddy 2011), and candidates were cross-validated using Phobius (Kall et al. 2007). Only proteins predicted to contain exactly seven transmembrane helices were retained as GPCRs.

Trypsins were identified based on Pfam domain PF00089 and confirmed using the MEROPS database (family S01A)(Rawlings 2006; Mistry et al. 2021). Putative lactonases were identified by BLASTp searches against the UniProt Swiss-Prot database (The UniProt Consortium et al. 2025), retaining the top hit per query (E-value ≤ 1e-5). Enzyme Commission (EC) numbers were assigned based on Swiss-Prot accessions, and proteins annotated with EC numbers corresponding to lactonase activity were extracted for downstream analyses.

### Detection of positive selection and functional enrichment

Positively selected genes (PSGs) were identified using the branch-site model implemented in CODEML (PAML v4.10.7)(Yang 2007)., with foreground branches specified to test for episodic positive selection along targeted lineages. To assess lineage-specific shifts in selection intensity between Mr23 and Mr2575, RELAX analyses were performed in HyPhy v4.5(Kosakovsky Pond et al. 2020); RELAX estimates a selection intensity parameter (K) that scales the distribution of ω values on the test branch relative to background branches (K > 1, intensified; K < 1, relaxed), with significance assessed by a likelihood-ratio test(Wertheim et al. 2015). GO enrichment of PSGs was conducted using InterProScan-derived(Jones et al. 2014) GO annotations and topGO in R, with strain-specific PSG sets tested against the strain background of GO-annotated genes using Fisher’s exact test (“classic” algorithm); terms with P ≤ 0.05 were retained.

## Data availability

The genome sequences reported in this paper have been deposited in the National Center for Biotechnology Information (NCBI) database BioSample accessions SAMN54551730 (*Metarhizium robertsii* ARSEF 1046), SAMN54551731 (*Metarhizium robertsii* ARSEF 1120), SAMN54551729 (*Metarhizium robertsii* ARSEF 727), SAMN54551733 (*Metarhizium robertsii* ARSEF 1878), SAMN54551732 (*Metarhizium robertsii* ARSEF 2547), SAMN54551735 (*Metarhizium robertsii* ARSEF 2575), and SAMN54551734 (*Metarhizium robertsii* Mr1-16). All data are available in the main text or the supplementary materials.

## Acknowledgments

We thank Novogene for genome-sequencing services. Figures were generated using Biorender.

