## Supplementary Figures S1-S21 for "Ecological diversification without genomic reorganization: repeat dynamics and regulatory evolution in *Metarhizium robertsii*"

**This PDF file includes:**

Figs. S1 to S21

References

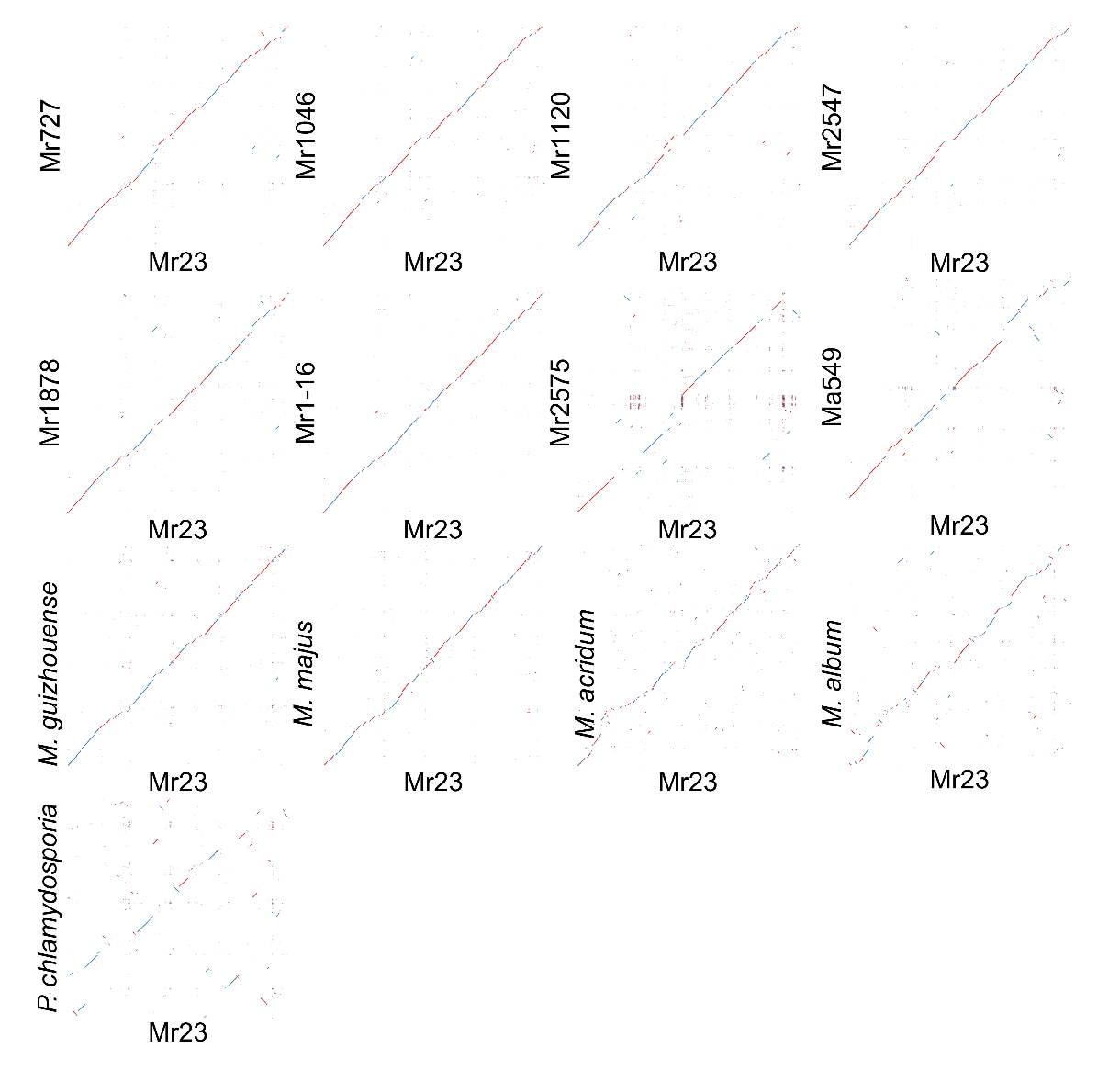

**Fig. S1. Whole-genome synteny comparisons between *M. robertsii* strains.**
Dot plots show pairwise genome alignments between Mr23 and other *M. robertsii* strains and related fungi, generated using NUCmer in the MUMmer v4.0.0 package. Scaffolds from Mr23 are plotted on the x-axis, and scaffolds from the comparison genome are shown on the y-axis. Diagonal alignments indicate regions of sequence similarity. Blue lines represent forward-strand alignments and red lines represent reverse-strand alignments.

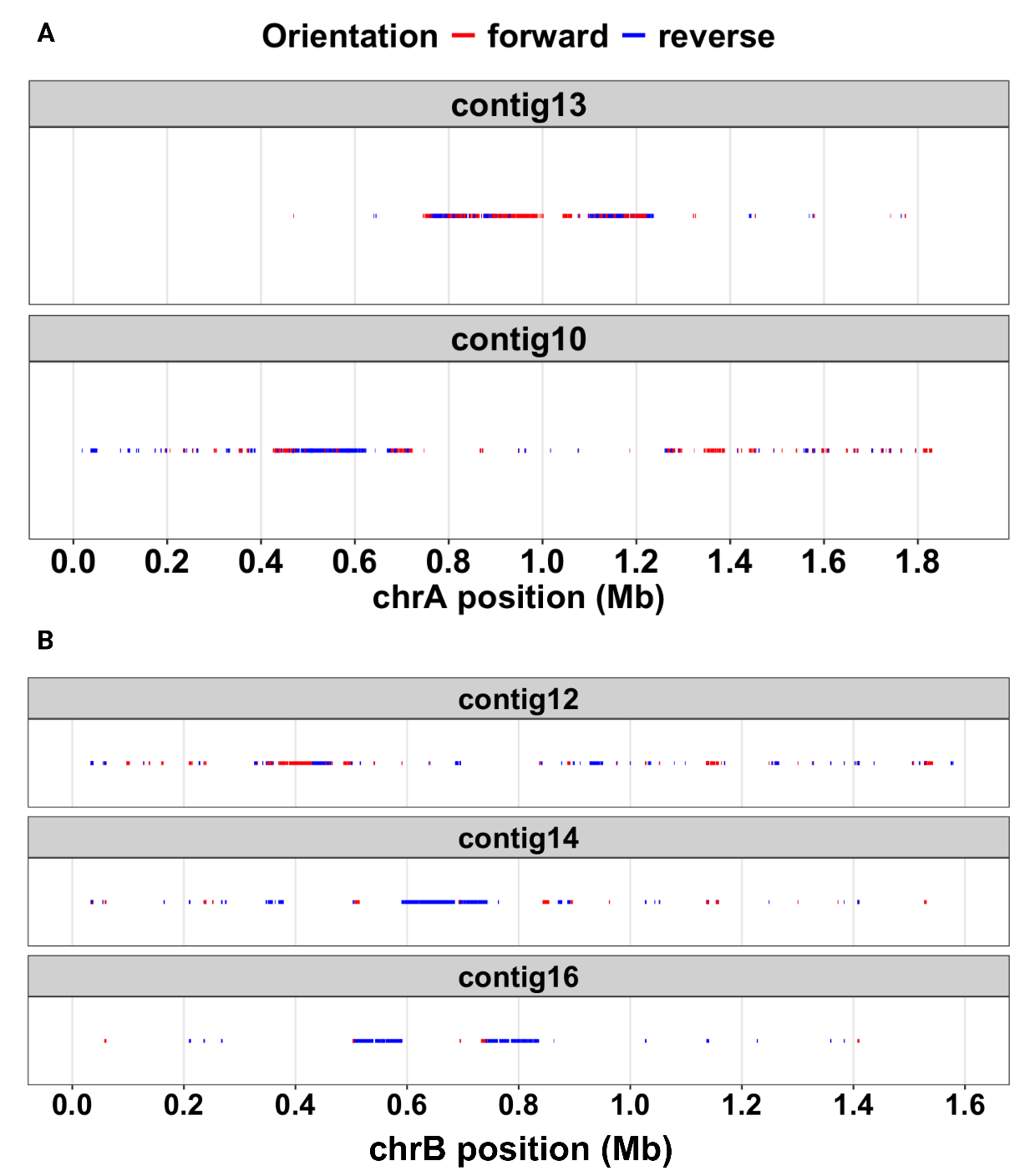

**Fig. S2. Distribution of chrA- and chrB-derived homologous sequences across PacBio contigs in Mr2575.**
(A) Alignment positions of chrA-derived sequences (from *M. robertsii* strain R3-I4) mapped onto contigs 10 and 13. (B) Alignment positions of chrB-derived sequences mapped onto contigs 12, 14, and 16. Each point represents a locally collinear alignment block, plotted by position along the reference chromosome (x-axis). Red and blue colors indicate forward and reverse strand alignments, respectively. The fragmented and dispersed distribution of alignment blocks across multiple contigs indicates that accessory chromosome sequences are not maintained as intact chromosomes but are instead integrated and rearranged within the genome.

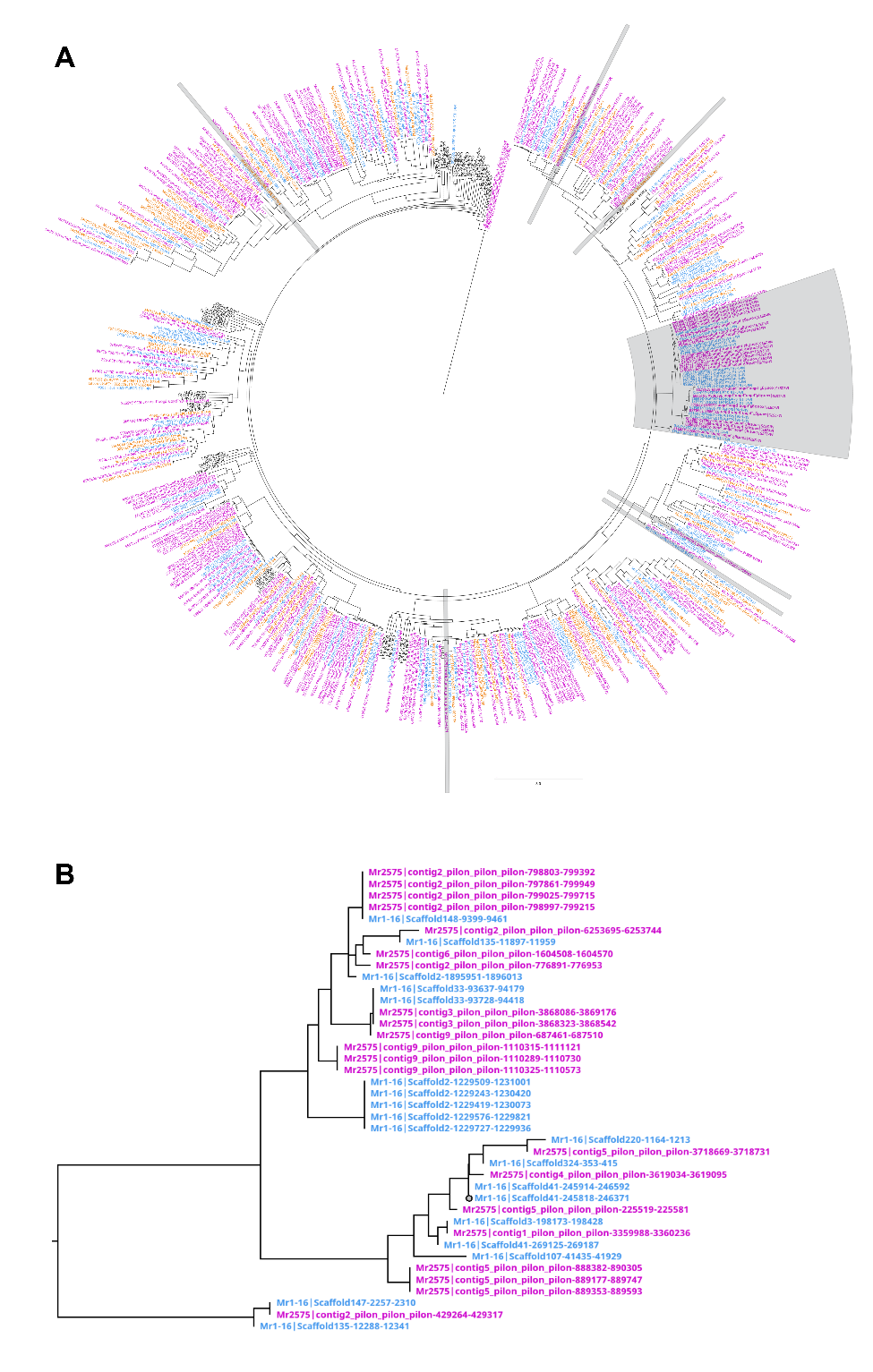

**Fig. S3. Phylogenetic relationships of LTR/Copia transposable elements in *M. robertsii* strains.**
Maximum-likelihood phylogeny of LTR/Copia transposable element (TE) sequences from Mr1-16 (blue), Mr2575 (magenta), and Mr23 (orange). TE copies located within RIP-affected genomic regions are indicated in grey. A clade enriched for RIP-associated TE copies (B), is shown enlarged for clarity. Scale bar indicates substitutions per site.

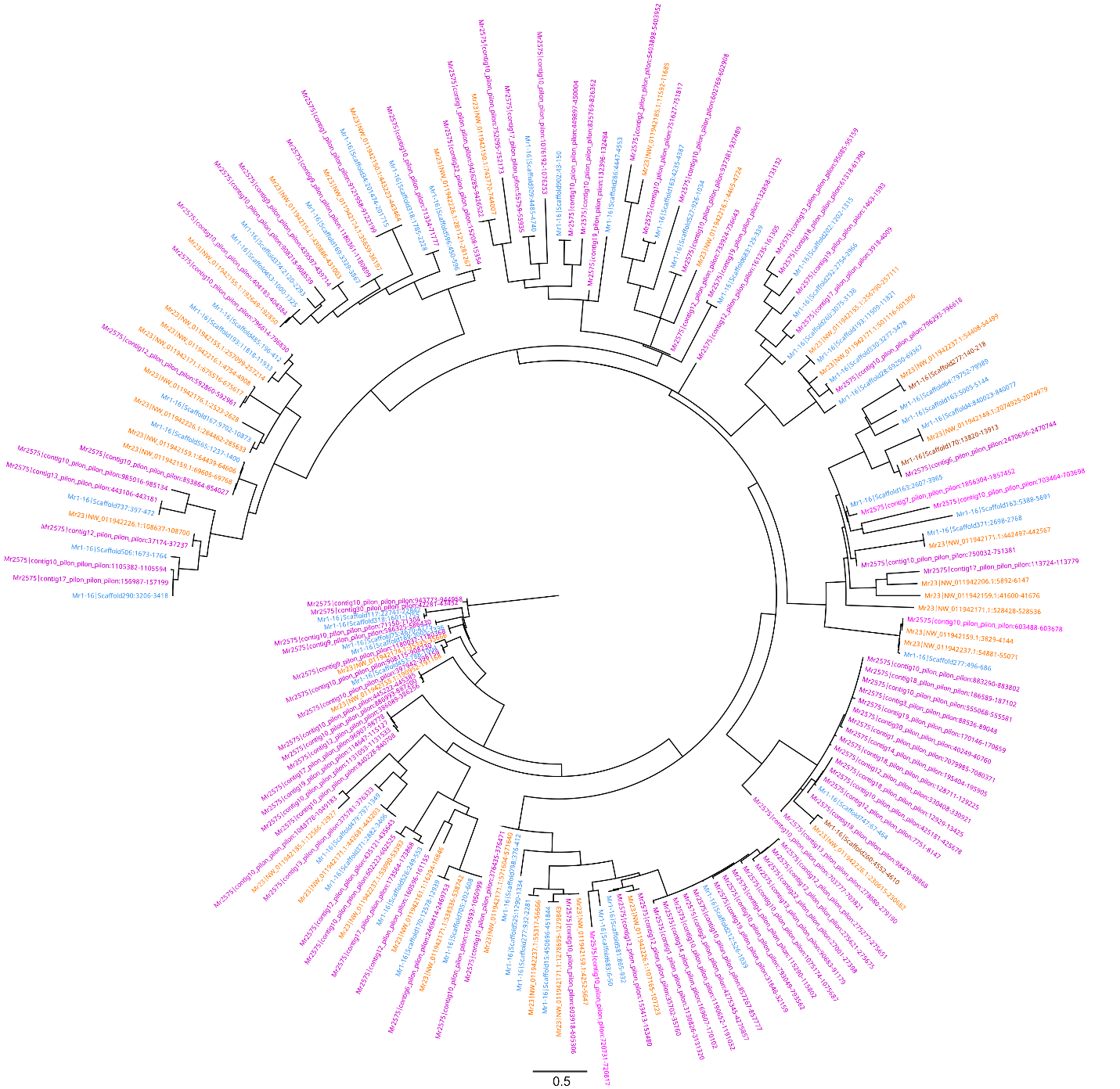

**Fig. S4. Phylogenetic relationships of DNA/hAT-Restless transposable elements in *M. robertsii* strains.**
Maximum-likelihood phylogeny of DNA/hAT-Restless transposable element (TE) copies from Mr1-16 (blue), Mr2575 (magenta), and Mr23 (orange). Scale bar indicates substitutions per site.

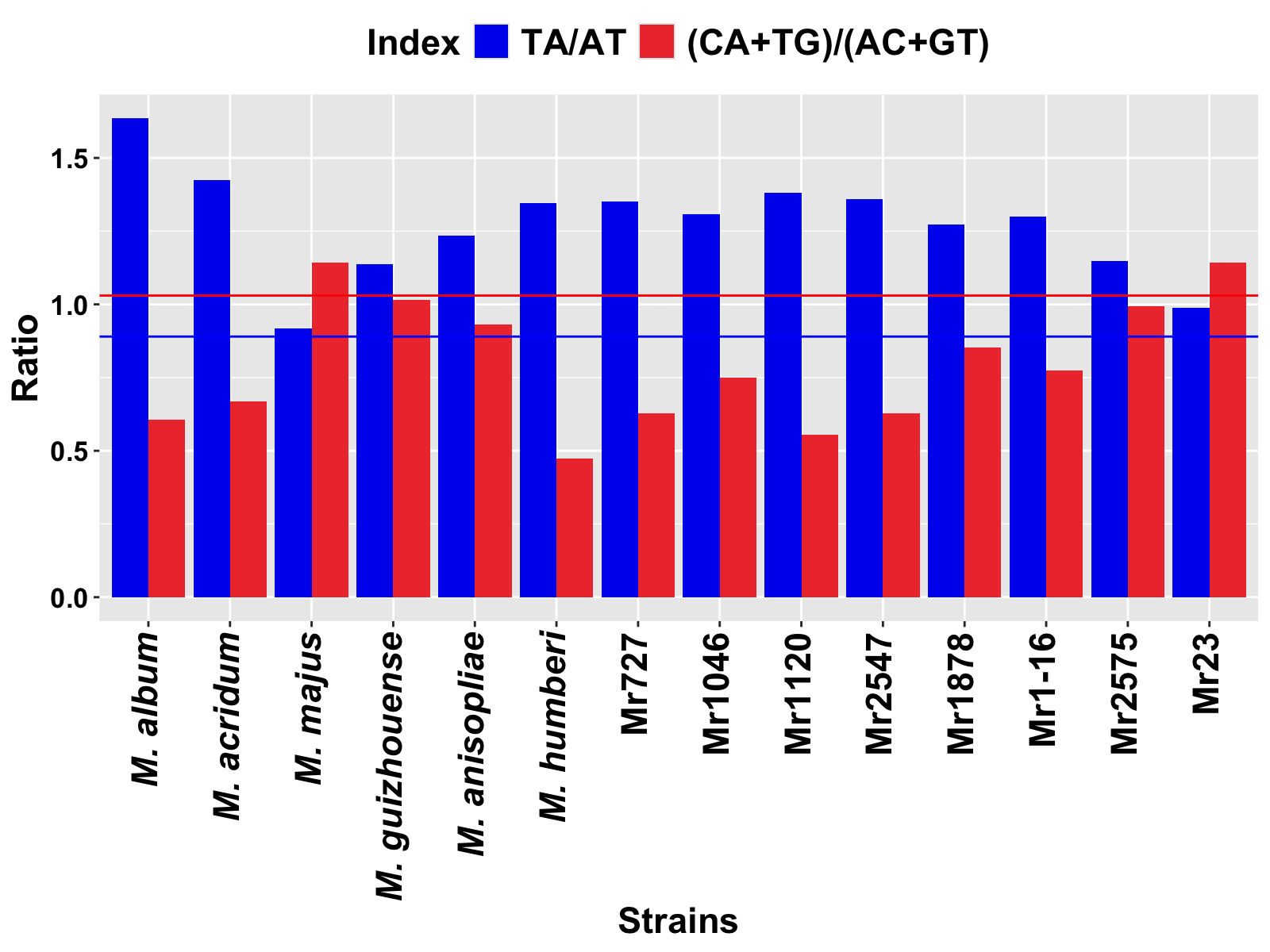

**Fig. S5. Analysis of Repeat-Induced Point (RIP) mutations in *M. robertsii* and other *Metarhizium* spp. using RIPCAL.** The RIP product (TA/AT) threshold is indicated by the blue line, and the RIP substrate [(CA+TG)/(AC+GT)] threshold by the red line. Strains with RIP product ≥ 0.89 and RIP substrate ≤ 1.03 are indicative of RIP in the genome(Gao et al. 2011).

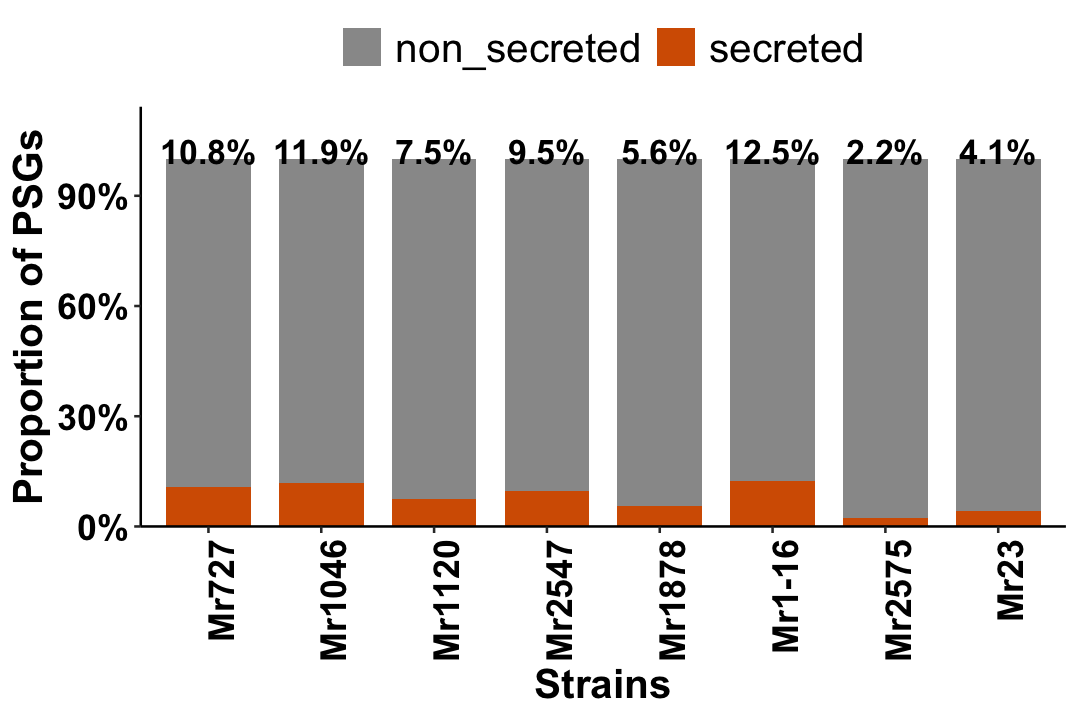

**Fig. S6.** **Proportion of secreted positively selected genes (PSGs) across M. robertsii strains.**
Stacked bar plot showing the proportion of predicted secreted (orange) and non-secreted (gray) PSGs in each *M. robertsii* strain. Bars are normalized to the total number of PSGs per strain. Across strains, only ~8% of PSGs are predicted to encode secreted proteins, indicating that positive selection is predominantly associated with intracellular functions rather than secreted effectors. Percentages above bars denote the fraction of PSGs predicted to contain signal peptides.

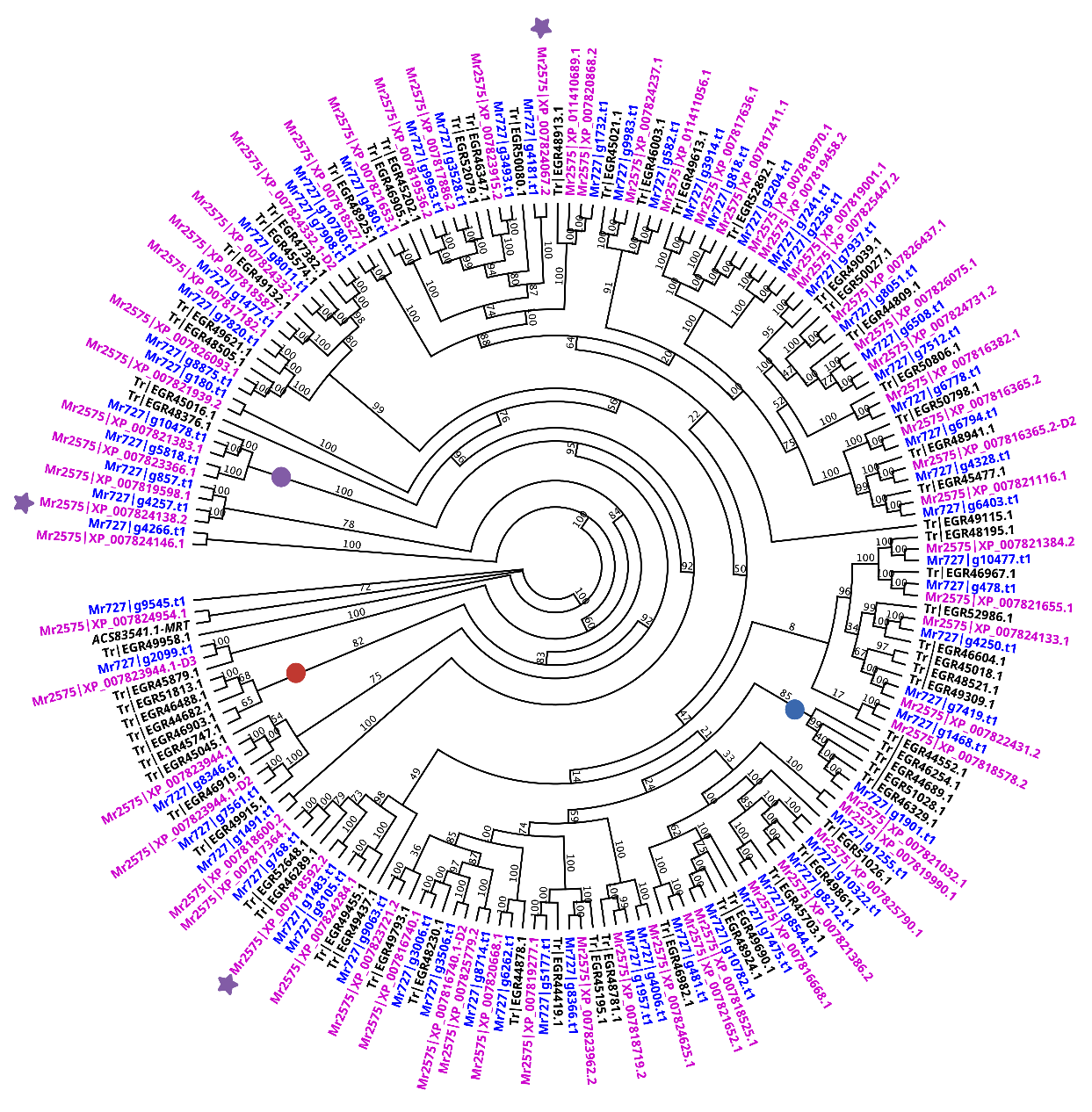

**Fig. S7. Phylogenetic analysis of sugar and other transporters (PF00083) in *M. robertsii* (Mr2575, Mr727) and *Trichoderma reesei*.**

Maximum-likelihood phylogeny of PF00083 family transporters from two *M. robertsii* strains (Mr2575 and Mr727) and *T. reesei*. The oligosaccharide transporter (MRT) from Mr23 is indicated in italics. A clade containing six *T. reesei* transporters annotated as maltose permeases is marked with a red circle. *T. reesei* encodes four quinate permease–like transporters, whereas Mr727 and Mr2575 each encode a single homolog (blue circle). A clade enriched for trehalose-associated transporters is indicated (purple circle). Purple stars denote three Mr2575 positively selected genes (PSGs).

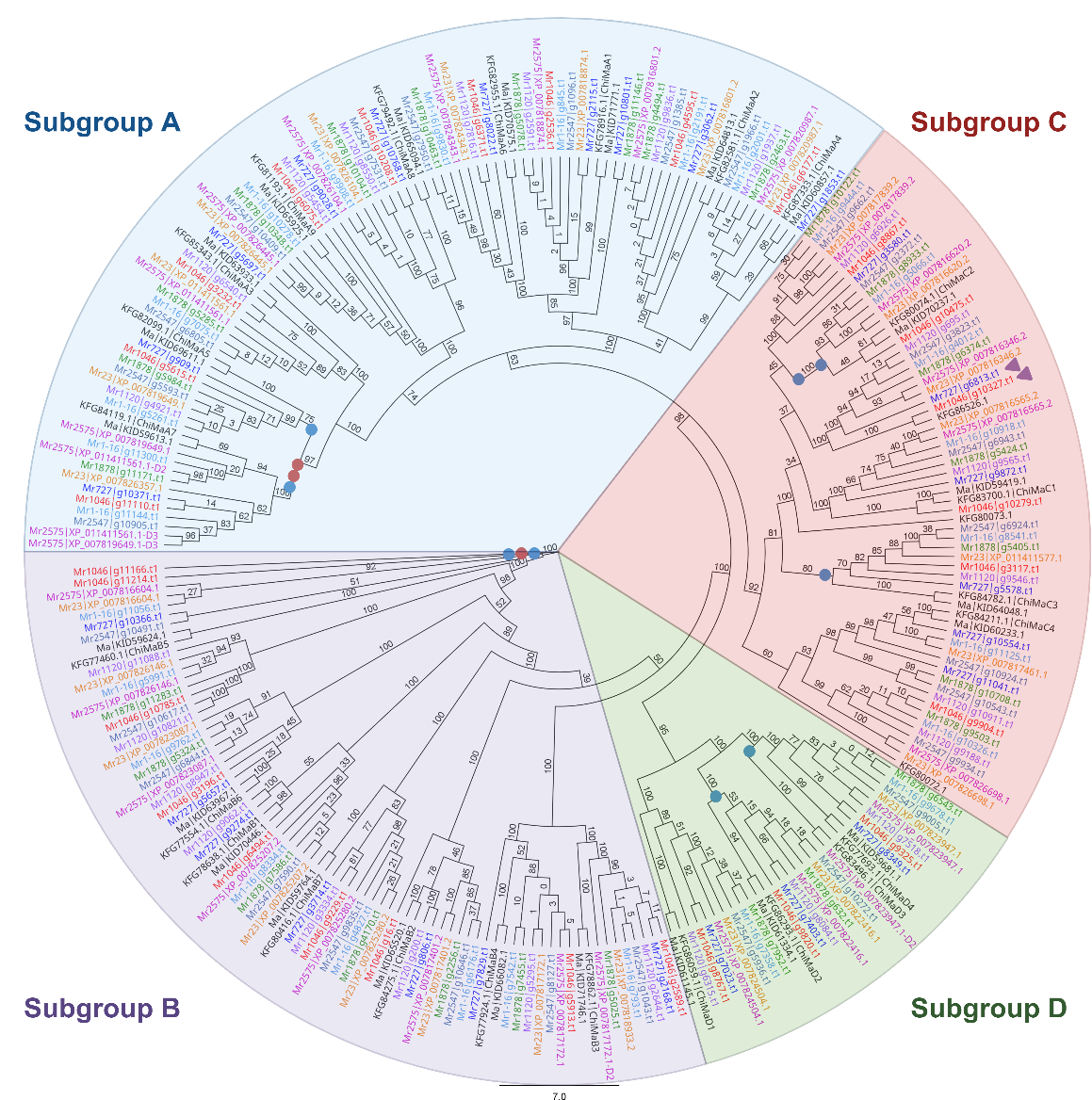

**Fig. S8. Phylogenetic relationships and lineage-specific turnover of GH18 chitinases in *M. robertsii*.**

Maximum-likelihood phylogeny of GH18 chitinases from eight *M. robertsii* strains and *M. anisopliae* Ma549. Conserved subfamilies shared across *M. robertsii* and *M. anisoplia*e strains (Ma549, and E6) include A2, A3, A4, A6, A8, A9, B1, B2, B3, B4, B7, D1, and D4. Red circles indicate inferred gene duplication events and blue circles indicate inferred gene losses mapped onto the species tree using parsimony. Three duplication events were detected within the *M. robertsii* lineage, including one B5 duplication in Mr1046 and two A7 duplications in Mr2575. Multiple lineage-specific losses were observed across strains, including absence of A5 and C3 in Mr2575, A7, C2, and B5 in Mr1120, C2 in Mr727, B5 in Mr1878, and D2 in Mr1-16. The D3 chitinase is unique to Mr2575 among *M. robertsii* strains but is present in *M. anisopliae* E6, indicating intra-specific variation across both species. Purple triangles denote positively selected genes; positive selection was detected in C1 chitinases of Mr1046 and Mr727, although only the Mr727 sequence is predicted to be secreted. Scale bar indicates amino acid substitutions per site.

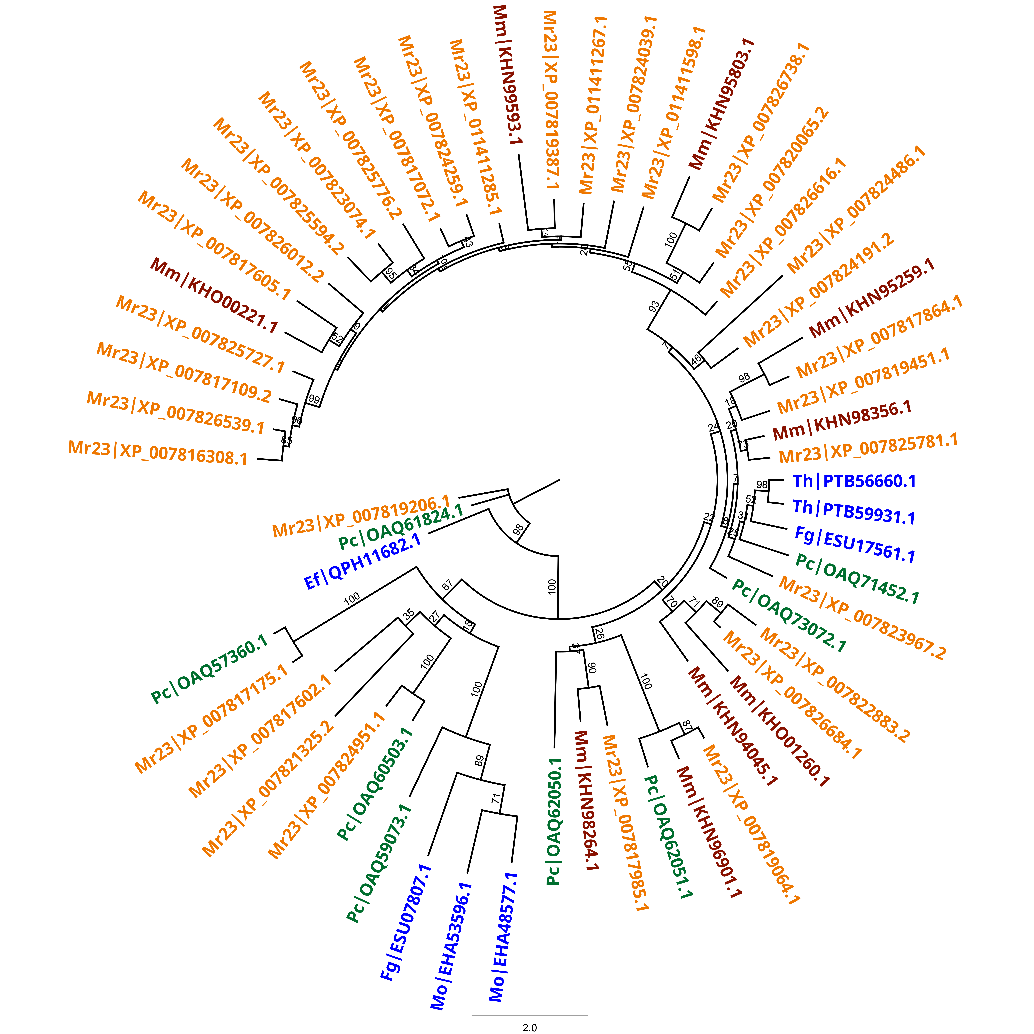

**Fig. S9. Phylogenetic relationships and expansion history of trypsins in *Metarhizium* and selected plant-associated fungi.**
Maximum-likelihood phylogeny of trypsin proteases from *M. robertsii* (Mr23), *M. album*, and the plant-associated species *Epichloë festucae*, *Trichoderma harzianum*, *Fusarium graminearum*, *Magnaporthe oryzae*, and *Pochonia chlamydosporia*. Bootstrap support values are shown for major nodes. Although several deep basal nodes exhibit low support, internal subclades are strongly supported. The Mr23 genome encodes 34 trypsins, *M. album* nine, *P. chlamydosporia* eight, *F. graminearum* two, *T.* *harzianum* two, *M. oryzae* two and *E. festucae* one. Six *P. chlamydosporia* trypsins cluster with Mr23 homologs, with only two retained in *M. album*, consistent with ancestral duplications predating divergence from *P. chlamydosporia* followed by gene loss in *M. album*. Five trypsins shared between Mr23 and *M. album* likely duplicated after divergence from *P. chlamydosporia*. The majority of additional Mr23 trypsins appear to have arisen through lineage-specific duplications after divergence from *M. album*. Green asterisks denote Mr23 trypsins previously reported to be highly expressed during insect cuticle penetration (Gao et al. 2011). The scale bar indicates amino acid substitutions per site.

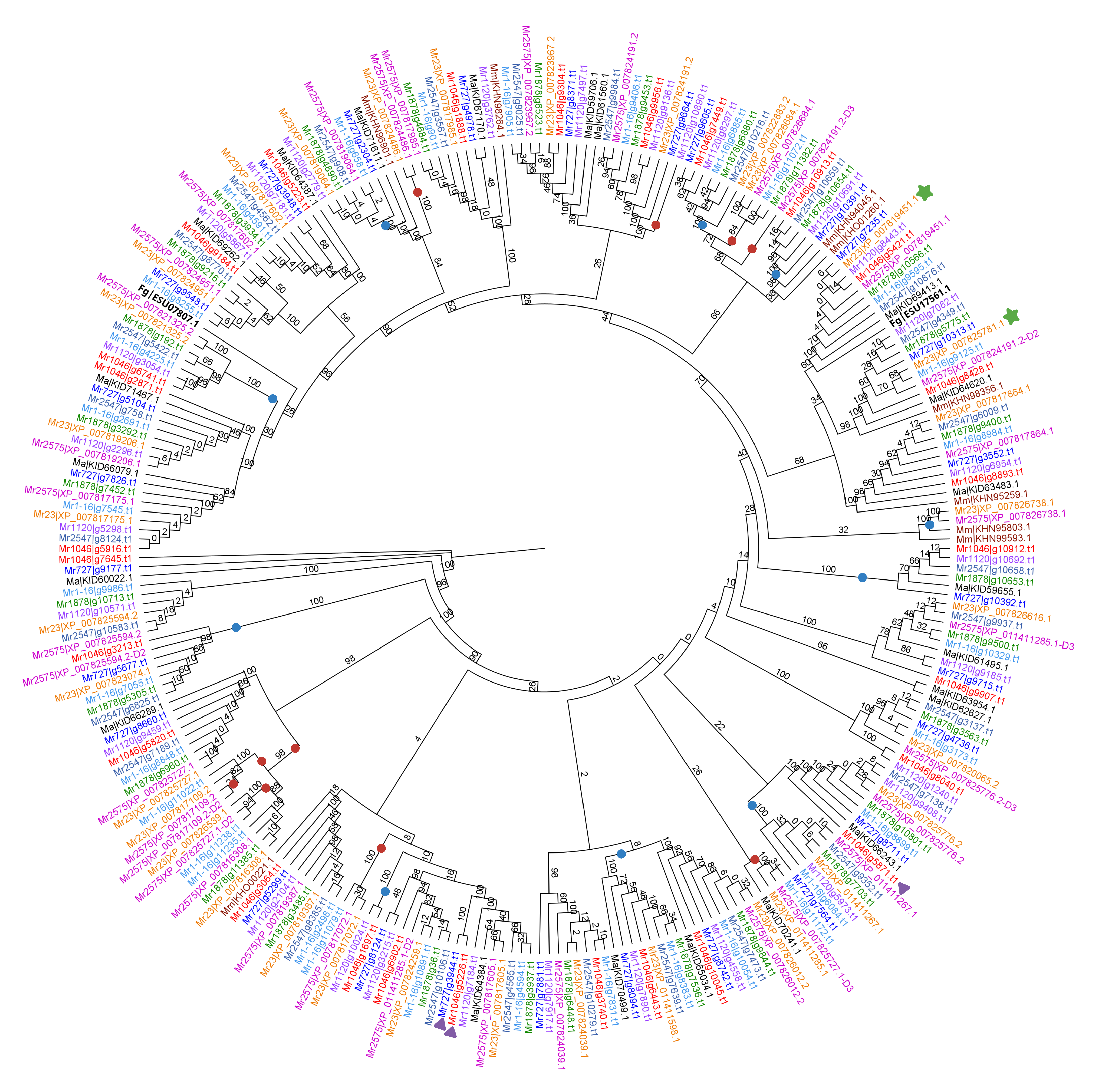

**Fig. S10. Phylogenetic analysis of trypsins in M. robertsii strains, M. anisopliae, M. album, and F. graminearum.**

Maximum-likelihood phylogeny of trypsins from eight *M. robertsii* strains, with *M. anisopliae* Ma549, *M. album*, and *F. graminearum* included as references. Single gene duplication events are indicated by red circles and inferred gene losses by blue circles, mapped according to parsimony onto the species tree. Ten gene loss events were detected across the phylogeny, including losses in Mr1120, Mr727, Mr1046 (two), Mr2575 (two), and in multiple ancestral nodes. Ten gene duplications were also inferred, including lineage-specific expansions in Mr2575 and in the common ancestors of Mr1-16, Mr23, and Mr2575. Purple triangles indicate genes identified as positively selected using branch-site analysis. Purple asterisks denote trypsins highly expressed during insect cuticle infection(Gao et al. 2011). One positively selected Mr1046 trypsin clusters with a positively selected Mr727 homolog (indicated by green stars). Two Mr23 trypsins highly expressed during infection form a conserved clade with one of the two *F. graminearum* trypsins, separate from duplication and selection events, suggesting functional conservation across Hypocreales. Overall, the pattern supports early expansion of trypsins in *Metarhizium* followed by modest recent turnover (ten gains and ten losses) within *M. robertsii* over approximately 3.4 MY.

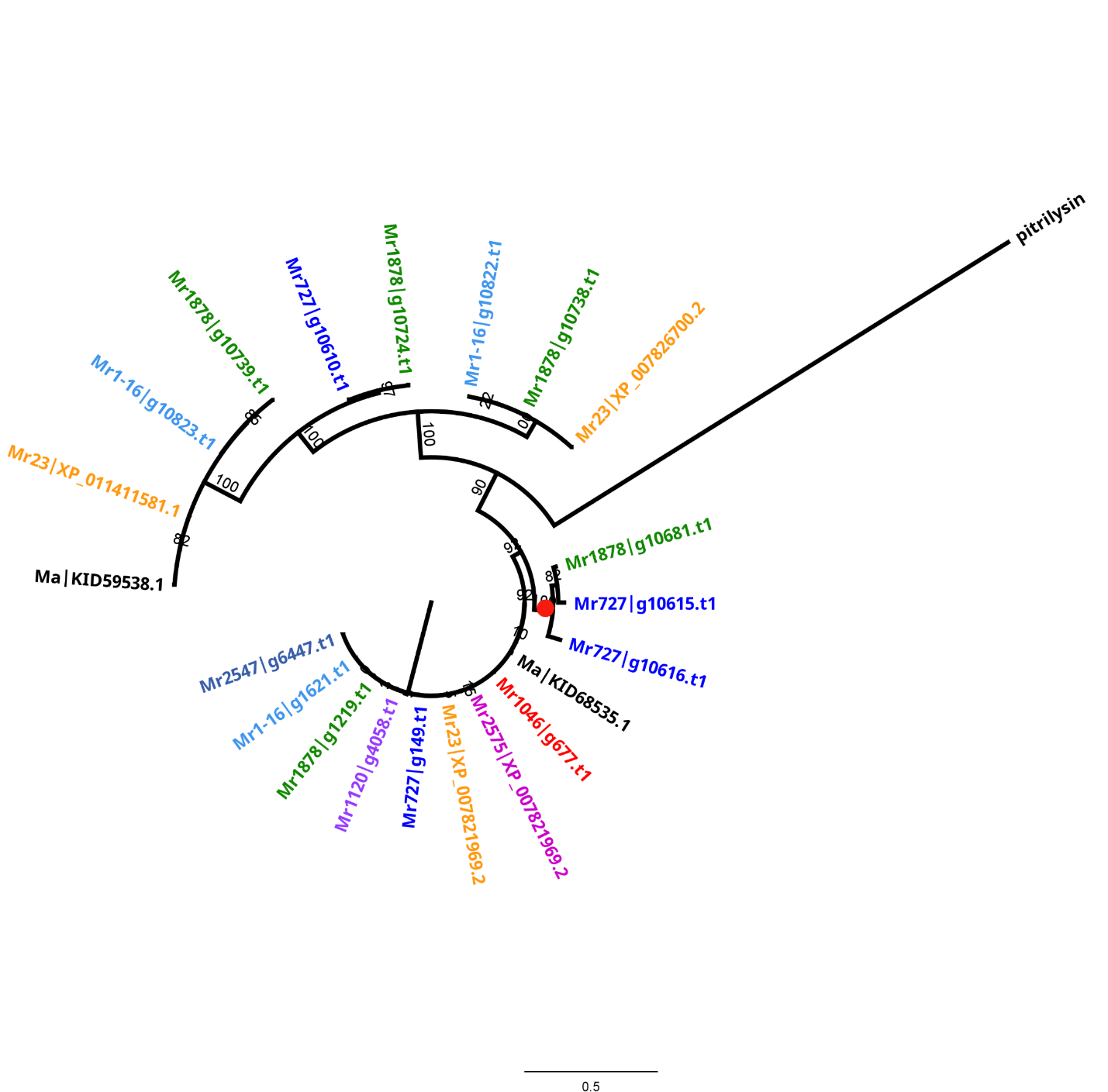

**Fig. S11. Lineage-specific expansion of M16A pitrilysin metalloproteases in *M. robertsii*.**

Maximum-likelihood phylogeny of M16A pitrilysin metalloproteases from eight *M. robertsii* strains, with *M. anisopliae* Ma549 included for reference. Most strains encode a single copy of the M16A pitrilysin implicated in a-factor mating pheromone production (Alper et al. 2006), whereas Mr1878 and Mr727 possess five and four copies, respectively. Red circles indicate an inferred recent gene duplication event since strain separation in Mr727. The scale bar indicates amino acid substitutions per site.

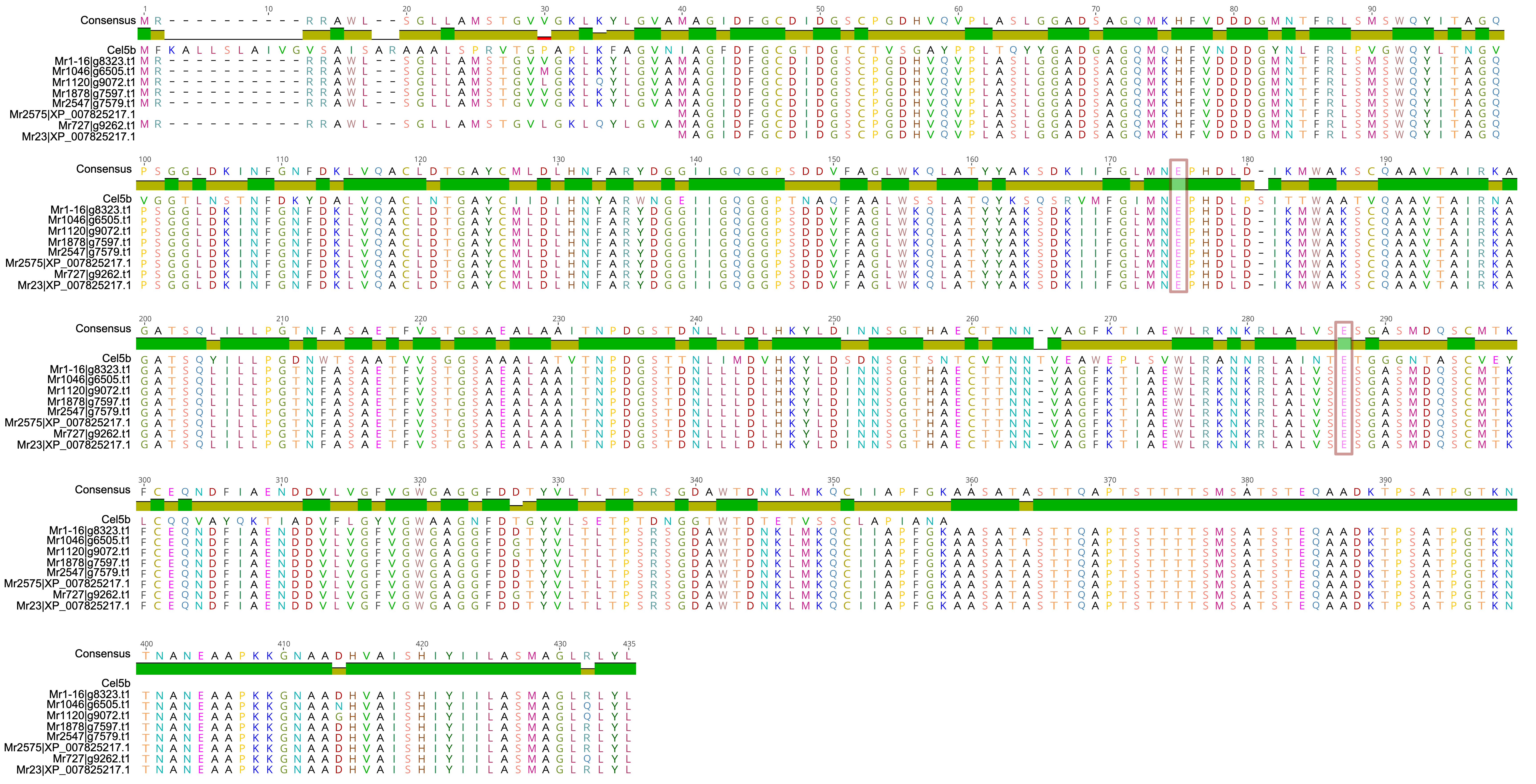

**Fig. S12. Sequence alignment of GH5_5 from *M. robertsii* and Cel5b from *Gloeophyllum trabeum*.**
Multiple sequence alignment comparing the GH5_5 protein from *M. robertsii* with Cel5b (NCBI accession XP_007867902.1) from the brown-rot fungus *G. trabeum*. The *M. robertsii* GH5_5 shares >49% amino acid similarity with Cel5b (E-value < 2.22 × 10⁻¹²⁶). Conserved catalytic glutamate residues (Glu175 and Glu287) are indicated and are shared between both sequences.

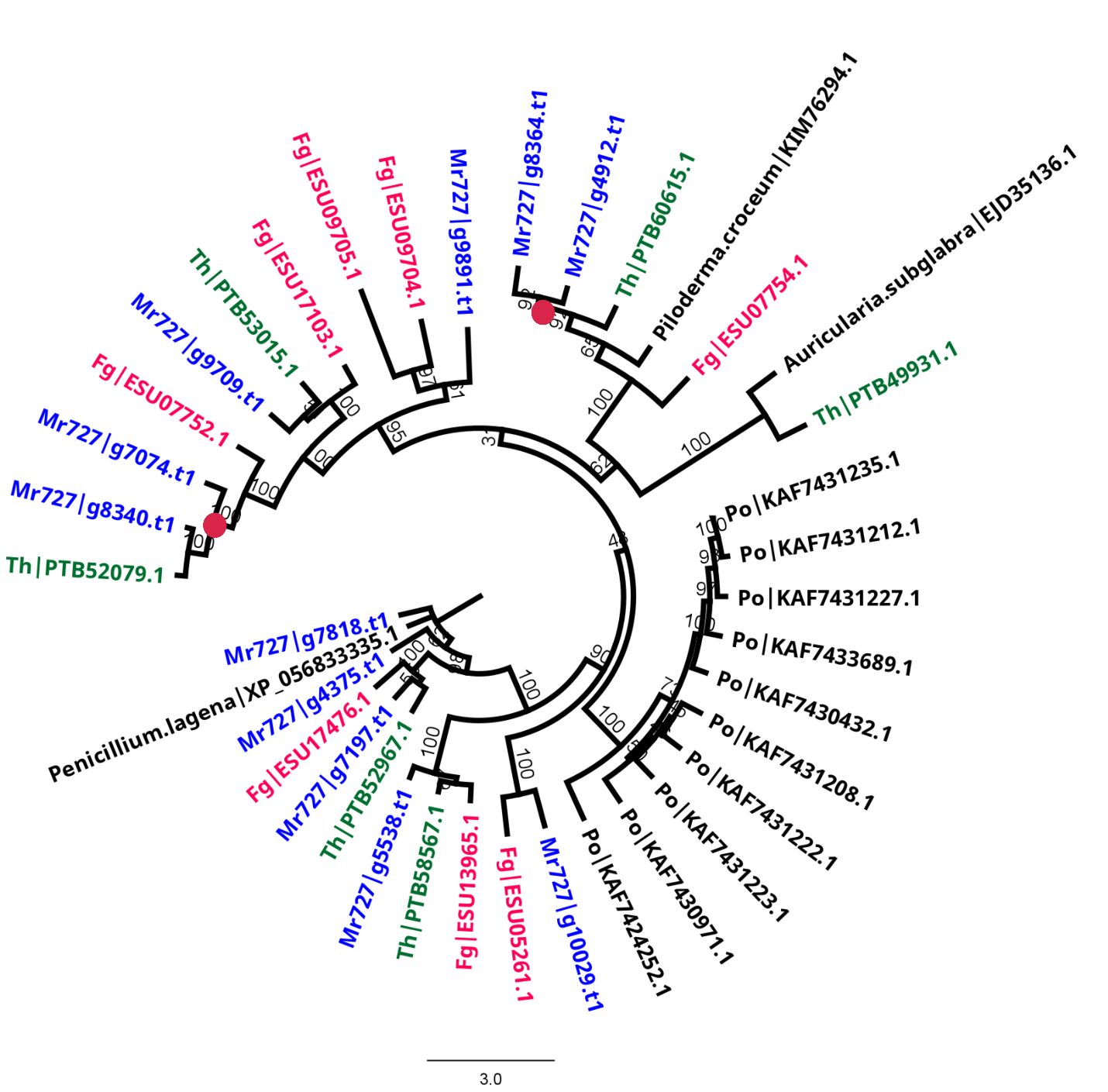

**Fig. S13. Phylogenetic relationships and lineage-specific expansion of AA1 laccases in *M. robertsii*.**
Maximum-likelihood phylogeny of AA1 multicopper oxidases (excluding AA1_2 subfamily members) from Mr727, *Trichoderma harzianum*, *Fusarium graminearum*, and *Pleurotus ostreatus*, a basidiomycete white-rot wood-decay fungus. Bootstrap values are shown for major nodes. Red circles indicate inferred single-gene duplication events within the *M. robertsii* lineage. The topology supports lineage-specific expansion of AA1 laccases in *M. robertsii*. Consistent with this pattern, *M. robertsii* encodes six to eight AA1_3 laccase-like multicopper oxidases, compared to one in *P. chlamydosporia* and two in *M. marquandii*. The scale bar indicates amino acid substitutions per site.

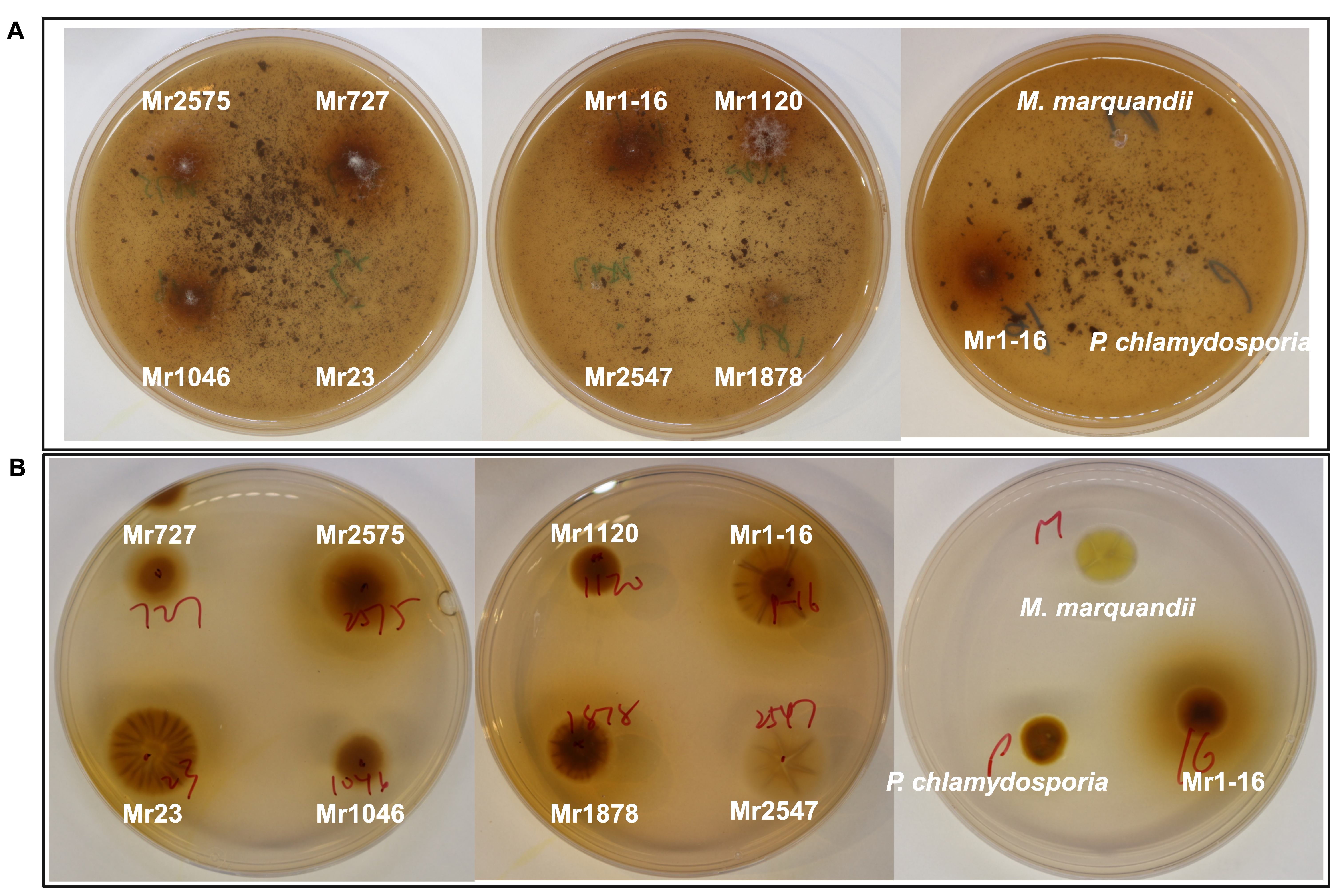

**Fig S14. Reverse views of biochemical plate assays for lignin- and tannin-associated activities.**
Reverse-side images of plates shown in Fig. 4. (A) Growth on minimal medium containing lignin as the sole carbon source. (B) Growth on PDA supplemented with 0.1% tannic acid. Images show *M. robertsii* strains, *P. chlamydosporia*, and *M. marquandii*.

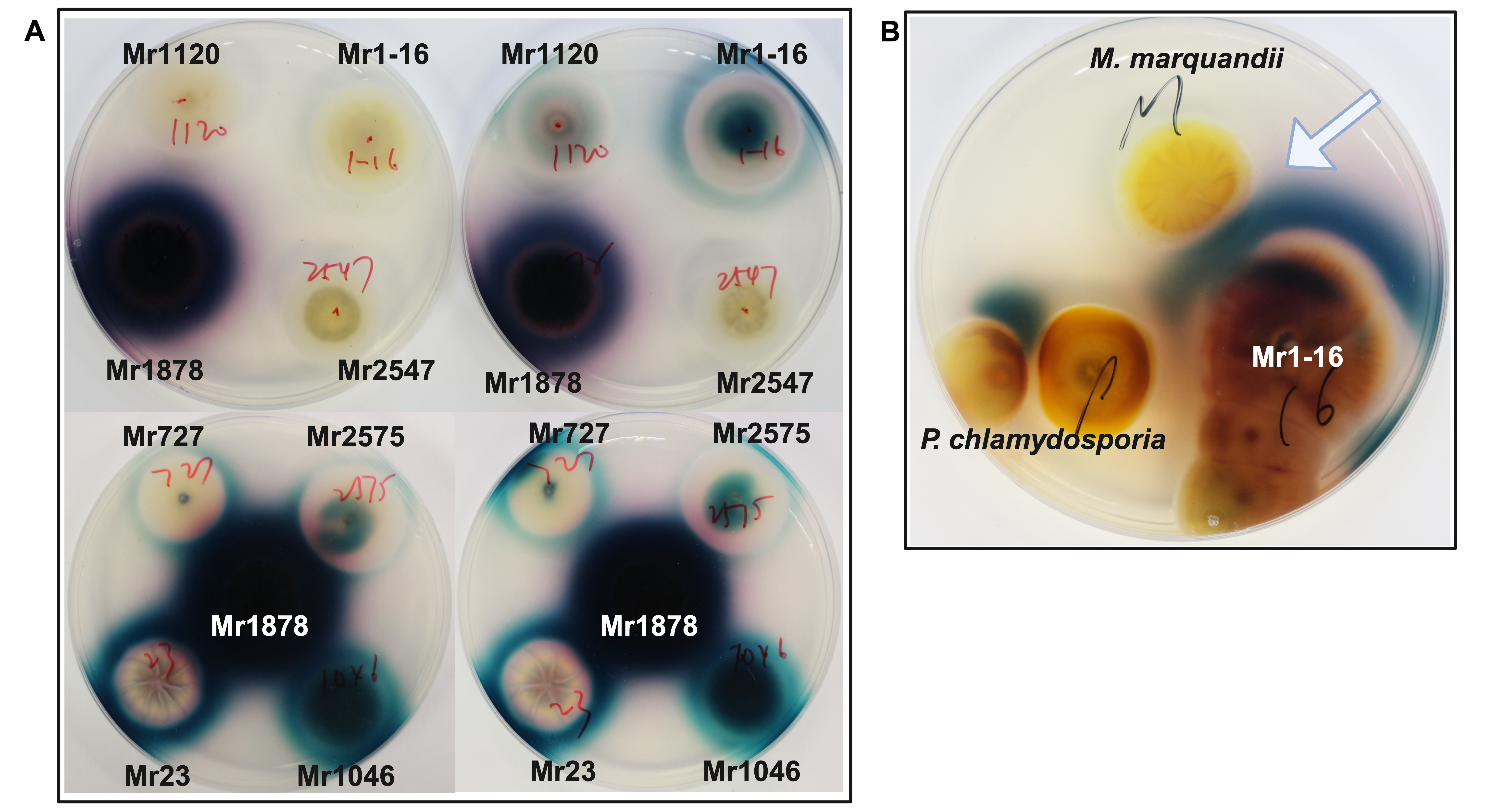

**Fig. S15. ABTS-associated oxidative activity in *M. robertsii* strains, *M. marquandii*,** **and *P. chlamydosporia*.**
(A) Colonies of Mr1878 rapidly produced large green halos on PDA supplemented with ABTS, which transitioned to blue over several days, indicative of high extracellular oxidative enzyme activity. In contrast, ABTS oxidation by the other strains was slower and variable, with oxidation products observed on some replicate plates but not others. (B) In cultures older than 7 days, Mr1-16 produced laccase activity localized to zones of interaction with *P. chlamydosporia* and *M. marquandii* (arrow).

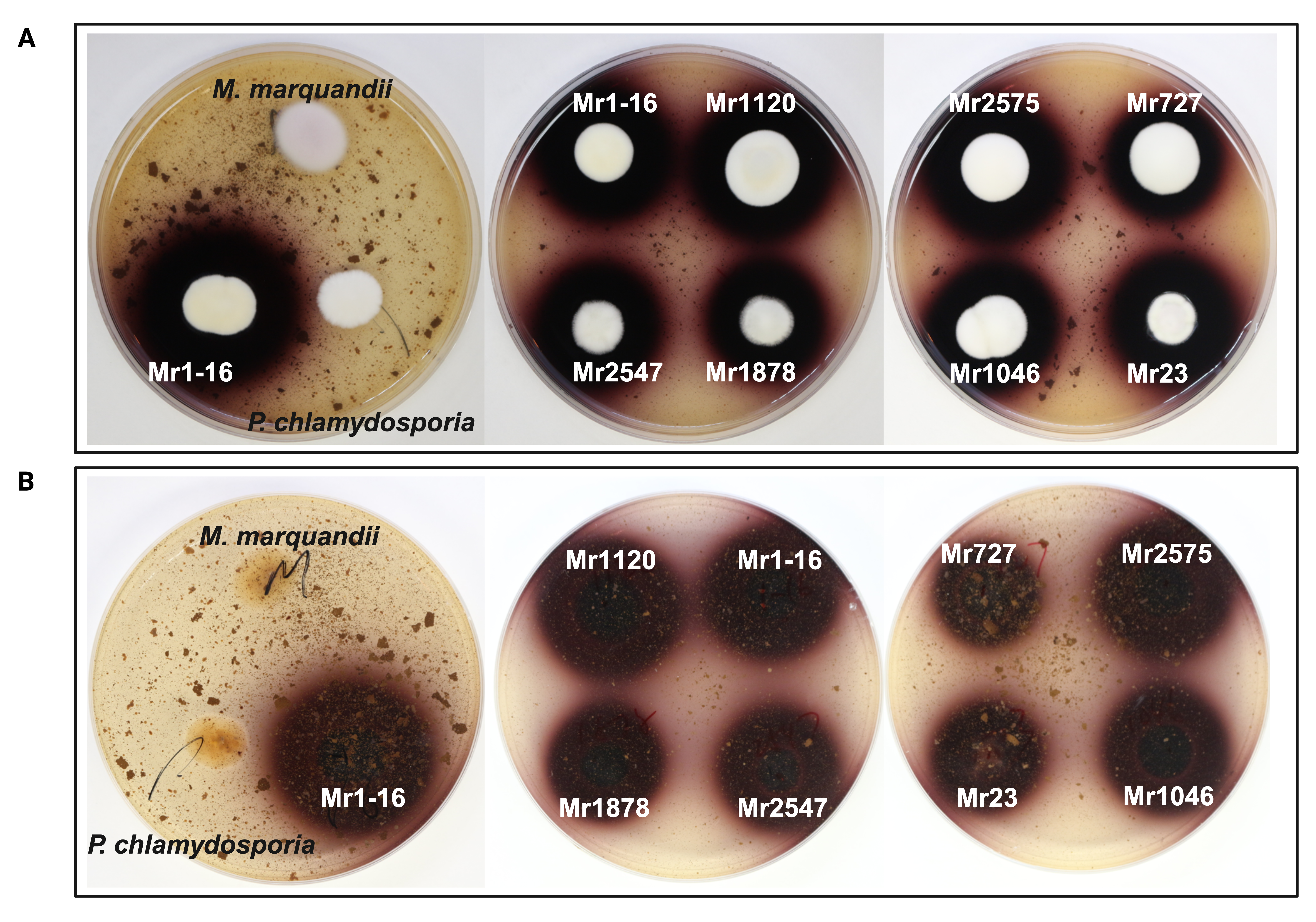

**Fig S16. Lignin- and ABTS-associated oxidative activity on PDA.**
Growth of *M. robertsii* strains, *P. chlamydosporia*, and *M. marquandii* on PDA supplemented with 1% lignin and ABTS. (A) Front view and (B) back view of colonies. All *M. robertsii* strains produced a purple halo, whereas *P. chlamydosporia* and *M. marquandii* showed trace or no discoloration.

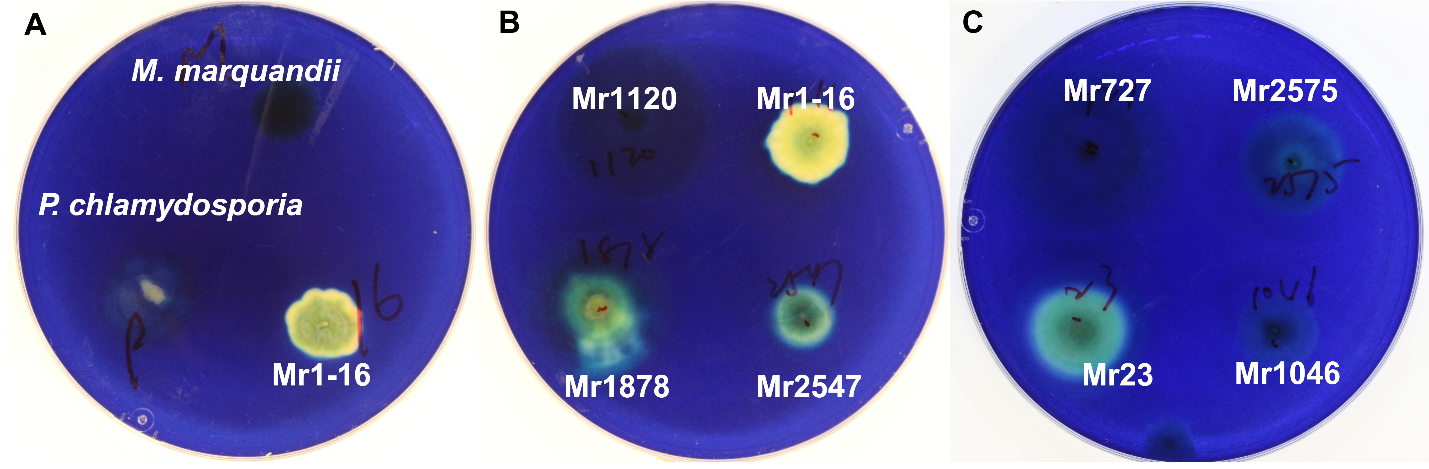

**Fig. S17. Methylene blue–associated redox activity on PDA.**
Growth of *M. robertsii* strains, *P. chlamydosporia*, and *M. marquandii* on PDA supplemented with methylene blue. (A) Partial decolorization beneath colonies of Mr1-16 indicates redox-mediated modification of methylene blue. In contrast, *P. chlamydosporia* and *M. marquandii* showed limited or no visible discoloration under the same conditions. (B) Localized decolorization was observed under colonies of Mr1-16, Mr1878, and Mr2547. (C) Localized decolorization was also observed under colonies of Mr23.

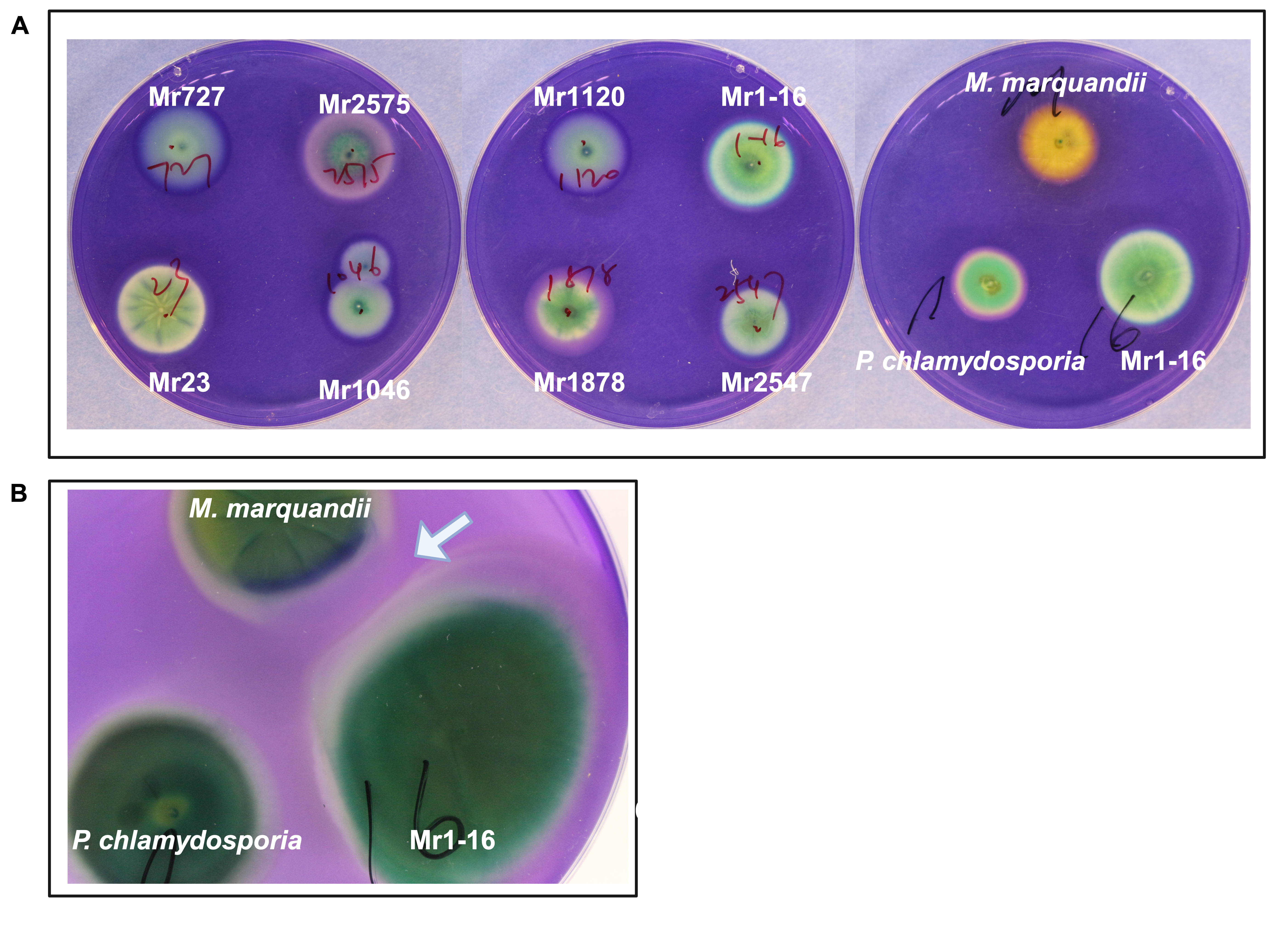

**Fig. S18. Azur B–associated oxidative activity on PDA.**
Growth of *M. robertsii* strains, *P. chlamydosporia*, and *M. marquandii* on PDA supplemented with azur B. (A) Partial decolorization, observed in *P. chlamydosporia* and *M. robertsii*, is indicative of azur B demethylation, whereas complete loss of color in *M. marquandii* indicates aromatic ring cleavage(Archibald 1992). (B) In older cultures, *M. robertsii* Mr1-16 showed complete decolorization beneath hyphae and at colony margins, particularly at confrontation zones with *M. marquandii* (arrow).

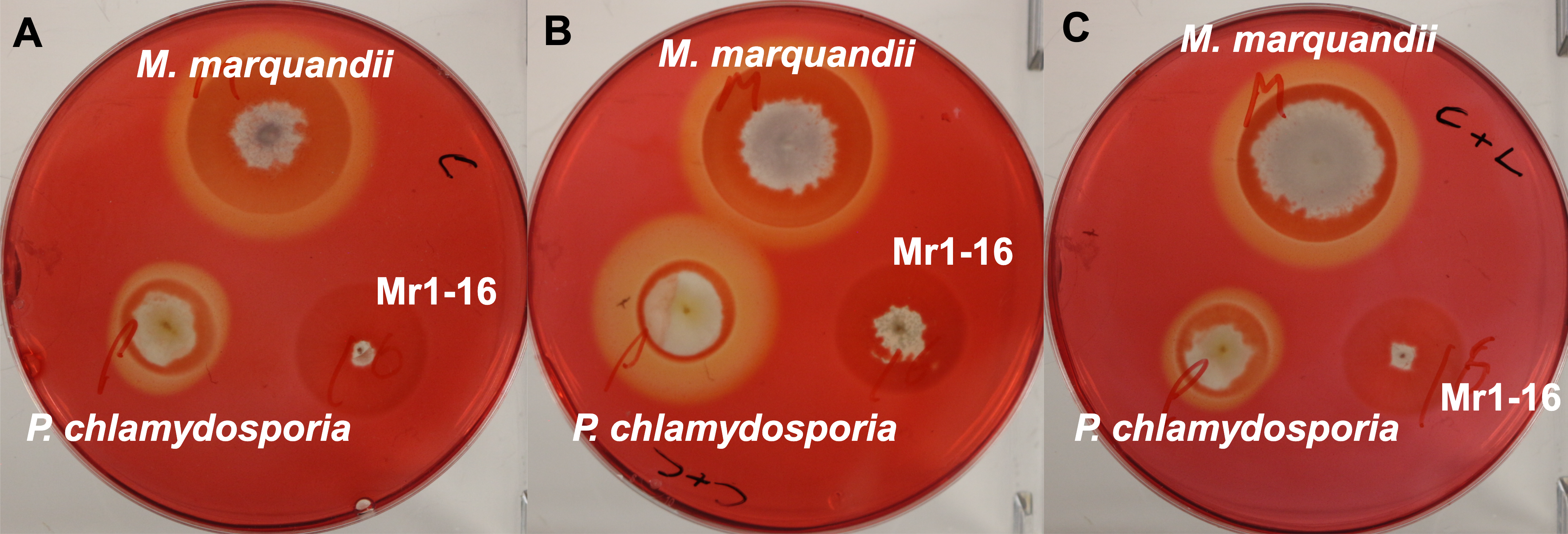

**Fig. S19. Cellulolytic activity of Mr1-16, *P. chlamydosporia*, and *M. marquandii* on carboxymethyl cellulose (CMC) with or without cellobiose or lactose.**
(A) Growth on agar plates containing CMC as the sole carbon source. *P. chlamydosporia* and *M. marquandii* produced large clear hydrolysis zones indicative of extracellular cellulase secretion, whereas Mr1-16 showed less radial growth and no detectable clearing beneath or surrounding colonies. (B and C) Effects of potential cellulase inducers on cellulase activity. Supplementation of CMC media with cellobiose (B) or lactose (C) did not alter colony size but differentially affected cellulase expression: cellobiose enhanced, whereas lactose repressed, cellulose hydrolysis by *P. chlamydosporia*. No cellulose degradation halos were detected for Mr1-16 and other *M. robertsii* strains (not shown) under any condition.

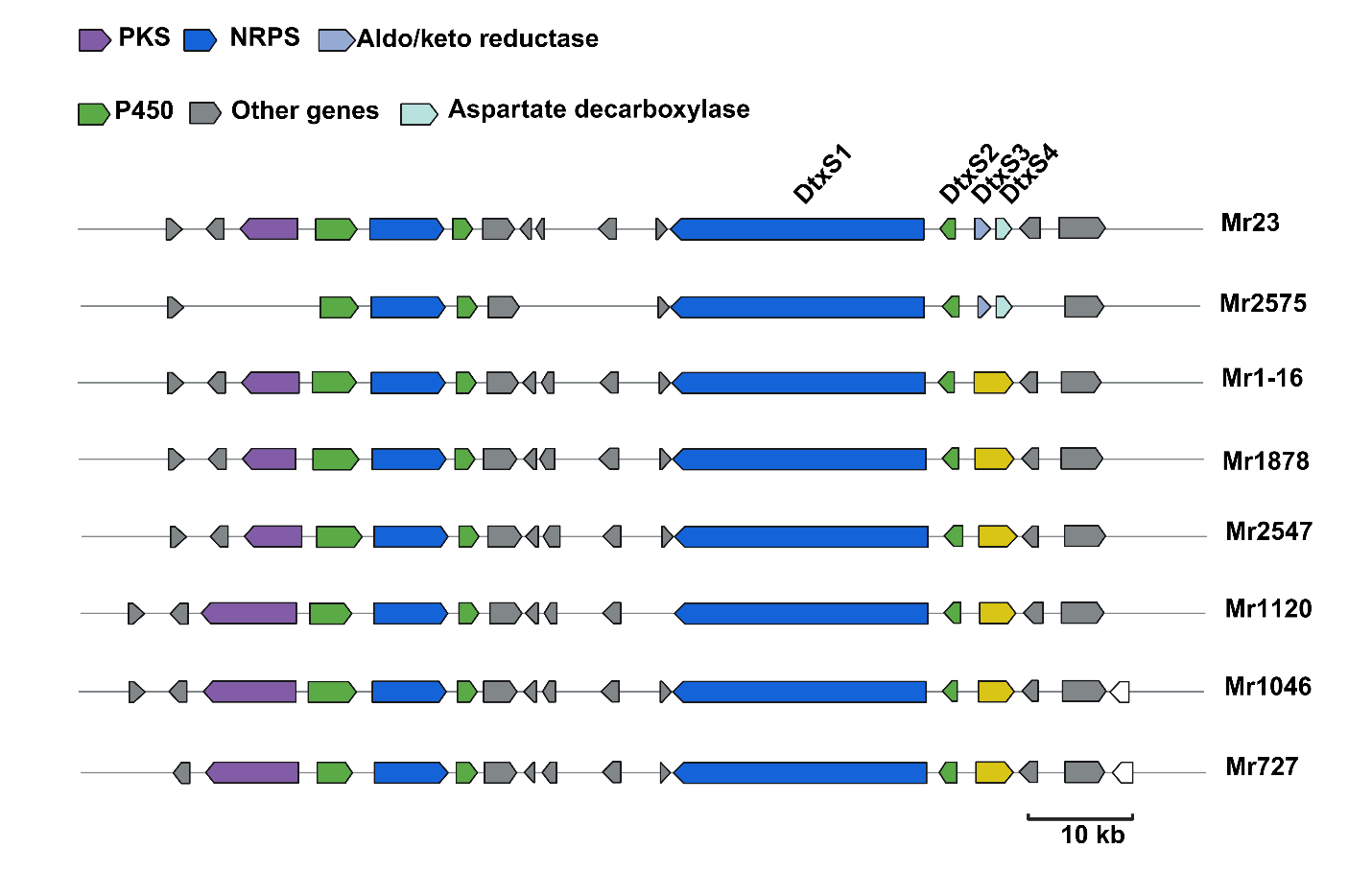

**Fig. S20. Organization and structural variation of the destruxin (dtx) biosynthetic gene cluster in *M. robertsii*.**
Schematic representation of the dtx gene cluster across *M. robertsii* strains. Core biosynthetic genes (dtxS1–dtxS4) are conserved across strains and share high sequence similarity (>98% for dtxS1 and dtxS2). Mr23 and Mr2575 retain adjacent dtxS3 and dtxS4 genes, whereas Mr727, Mr1046, Mr1-16, Mr1878, Mr2547, and Mr1120 encode a single fusion gene corresponding to dtxS3–dtxS4 (highlighted in yellow), with >96% amino acid similarity among strains. Mr2575 additionally lacks several upstream genes present in other strains. Overall gene content of the core biosynthetic locus remains conserved despite local structural variation.

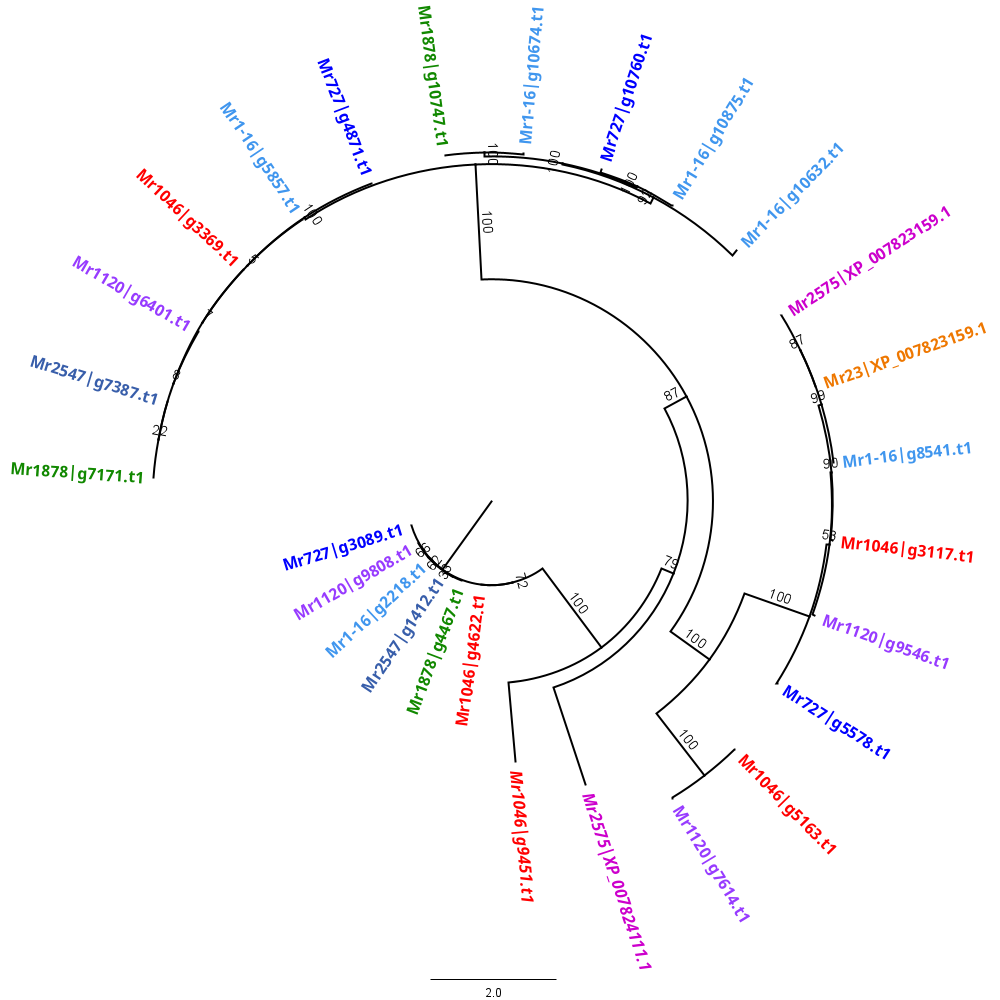

**Fig S21. Phylogenetic relationships of duplicated Zn2Cys6 transcription factors in *Metarhizium robertsii*.**
Maximum-likelihood phylogeny of duplicated Zn2Cys6 transcription factors (ZnFTFs) across eight *M. robertsii* strains. The scattered distribution of duplicated clades across the eight strains is consistent with ancestral duplication followed by gene losses in various strains. Duplicated genes arising after strain divergence shown in *italics*. Scale bar indicates substitutions per site.
